# A photosynthesis operon in the chloroplast genome drives speciation in evening primroses

**DOI:** 10.1101/2020.07.03.186627

**Authors:** Arkadiusz Zupok, Danijela Kozul, Mark Aurel Schöttler, Julia Niehörster, Frauke Garbsch, Karsten Liere, Irina Malinova, Ralph Bock, Stephan Greiner

## Abstract

Incompatibility between the cytoplasm and the nucleus is considered as major factor in species formation, but mechanistic understanding is poor. In evening primroses, a model plant for organelle genetics and population biology, hybrid offspring regularly displays chloroplast-nuclear incompatibility. These incompatibilities affect photosynthesis, a trait under selection in changing environments. Here we show that light-dependent misregulation of the plastid *psbB* operon (encoding core subunits of photosystem II and the cytochrome *b*_6_*f* complex), can lead to hybrid incompatibility, thus ultimately driving speciation. This misregulation results in an impaired light acclimation response in incompatible plants. Moreover, as a result of their different chloroplast genotypes, the parental lines differ in their photosynthesis performance upon exposure to different light conditions. Significantly, the incompatible chloroplast genome is naturally found in xeric habitats with high light intensities, whereas the compatible one is limited to mesic habitats. Consequently, our data raise the possibility that the hybridization barrier evolved as a result of adaptation to specific climatic conditions.

## Introduction

Incompatibility between nuclear and organellar genomes represents a mechanism of reproductive isolation observed in a wide range of taxa^1–8^. However, with the exception of the commercially important trait cytoplasmic male sterility (CMS), little is known about the molecular and evolutionary mechanisms of cytoplasmic incompatibility (CI). Mechanistic studies are available from only a handful of cases^1,3,9–12^, contrasting the great biological importance of the phenomenon. Increasing evidence accumulates that CI arises early in the separation of two genetic lineages^1,5,7,8,13,14^, and thus, represents an initial barrier towards reproduction isolation. This suggests that CI acts as a major factor in species formation, making a mechanistic understanding of its molecular basis and in a population genetic context highly desirable.

The evening primrose *(Oenothera)* is a plant model uniquely suited to address the mechanisms of reproductive isolation through hybrid incompatibility. Crosses between *Oenothera* species usually produce viable offspring that, however, regularly displays incompatibility between the chloroplast and the nuclear genomes (plastome-genome incompatibility; PGI). These incompatibilities represent the only strong hybridization barrier between *Oenothera* species, which often co-occur in overlapping ecological niches, within hybridization zones^8,15,16^ (Fig. 1). Hybrid incompatibility of nuclear loci is essentially absent^17,18^, underscoring the importance of CI as the cause of incipient isolation of the hybrid. In addition, *Oenothera* is a prime example for hybrid speciation^19,20^, in that permanent translocation heterozygosis, a form of cross-inducible functional asexuality, can occur. Such genotypes display a meiotic ring and bread true upon self-fertilization. In crosses, this can lead to an immediate fixation of a hybrid^21–23^ (also see Methods and below).

**Fig. 1.**
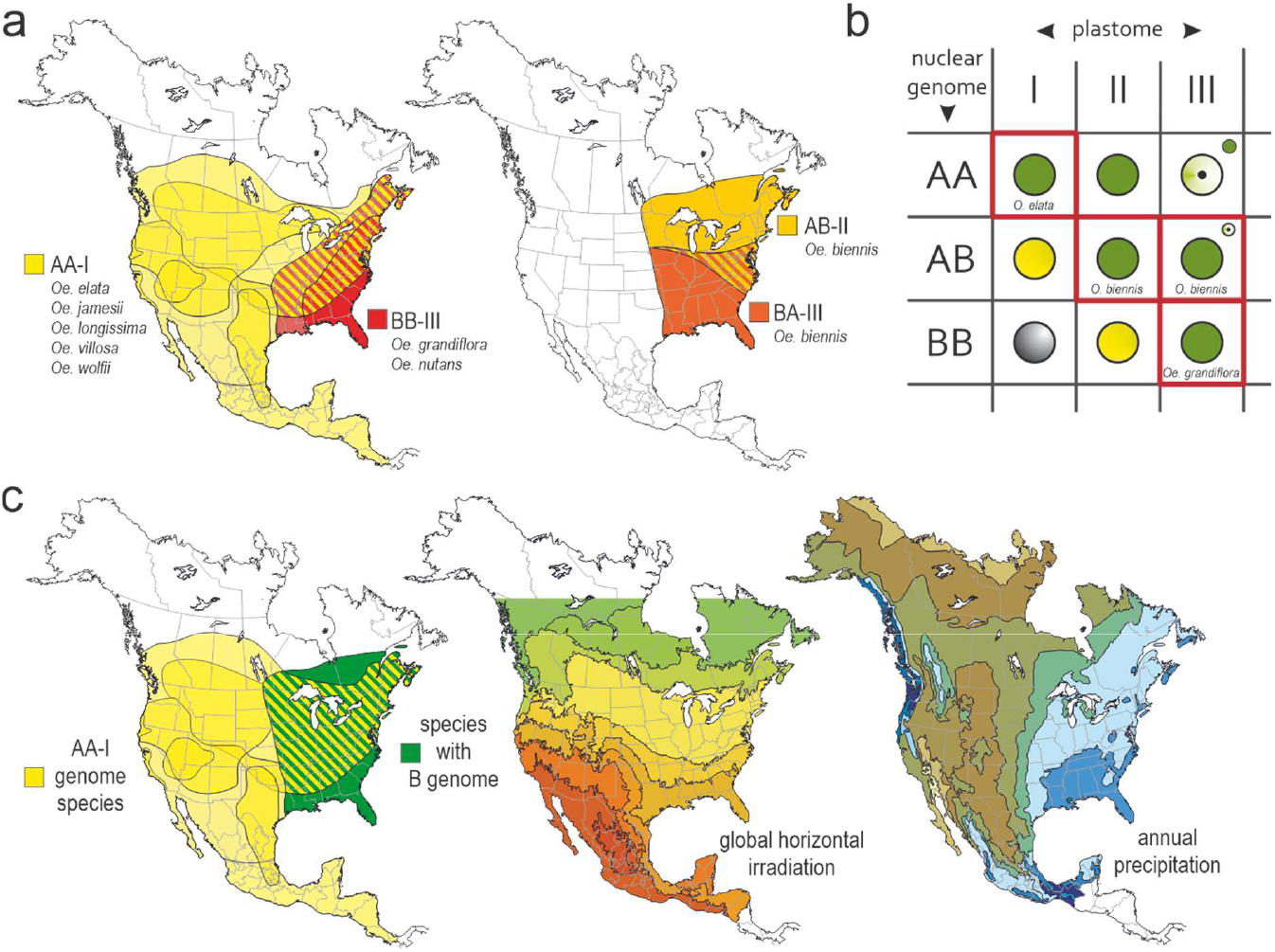
Distribution of *Oenothera* AA-I, BB-III and AB-II/BA-III species, and compatibility/incompatibility relations upon hybridization. **a**, Native distribution of *Oenothera* A and B genome species and their hybridization zones. **b**, Genetic species concept of evening primroses, based on plastome/nuclear genome compatibility and incompatibility, exemplified for the A and B genome species. Species are defined by their combinations of nuclear and chloroplast genomes (boxed in red), and are genetically separated by PGIs that occur upon hybridization and vary in the severity of the hybrid phenotype (BB-I white, AB-I and BB-II yellow-green, AA-III bleaching leaf phenotype). **c**, Association of AA-I and B genome (BB and AB) species of *Oenothera* to the xeric and mesic habitats of North and Central America. See text for details. Distribution maps redrawn from Dietrich *et al.* (1997)^16^. Climate data are from SolarGis and North American Environmental Atlas.

In evening primroses, three genetic lineages (A, B, and C) exist and are separated by PGI^8,15,21^. The genetic lineages occur as basic nuclear genome types in a homozygous (as AA, BB, or CC) or stable heterozygous (as AB, AC, or BC) state, and can be combined with five basic chloroplast genome types (I–V). The presence of distinct nuclear and chloroplast genomes, and the sexual separation of the species by PGI has led to the development of a genetic species concept for Oenothera^15,16,18,24^. Specific combinations of nuclear and chloroplast genomes define the species^16^. Other genome combinations can occur in weak or inviable hybrids, thus sexually separating the species (Fig. 1b). Strikingly, these chloroplast-mediated speciation barriers rely on photosynthesis, a trait under selection in changing environmental conditions^25,26^. This makes *Oenothera* an appealing model to understand the genetic basis of speciation. The incompatibility loci separating the species are relevant for speciation by definition and may even be a result of adaptive evolution.

For example, hybridization between *Oenothera elata* (an AA-I species) and *O. grandiflora* (a BB-III species) produces the incompatible combination AB-I. This genetic incompatibility poses the hybridization barrier between AA and BB species that appears to prevent colonization of western North America by B genome species (Fig. 1c)^8^. It should be emphasized that hybridization is frequent in the genus. As a rule, it occurs if plastome-genome combinations permit. Importantly, all viable combinations can be confirmed in hybrids in nature^16^. Hence, PGI appears to act as a major mechanism preventing gene flow. In addition, ecological species separation is likely to occur. For example, existence of the green and compatible hybrid AA-II can be confirmed, but it does not establish stable populations in nature^16^ (Fig 1a,b).

It is assumed that climate changes and periods of glaciation during the Pleistocene have shaped the genetic and ecological characteristics of the basic lineages A, B and C^21^. This is well supported by the estimated divergence time of the chloroplast genomes^27^ and nuclear genome variation^28,29^. Following this view, the three lineages originated from Middle America and reached the North American continent in several waves. The lineages resemble the ancestral sexual and homozygous species AA-I, BB-III and CC-V, and crosses between them usually result in PGI^8,15^. However, during the Pleistocene, hybridization between the basic lineages must have happened that produced viable offspring^16,20,21^. Those were fixed in the structural heterozygous and functional asexual species (AB-II or BA-III, AC-IV, and BC-IV). Hence, especially plastome II and IV can be seen as relict genotypes of earlier stages of plastome evolution^15,16^. Finally, it should be mentioned that also recent plastome divergence appears to be a consequence of glaciation. Separation of plastome II and III in the two major subpopulations of *O. biennis* coincides with the expansion of the Wisconsin glacier, suggesting post-glaciation dispersal events^30^ (Fig 1a).

The aim of this work was to understand the mechanism of the AB-I incompatibly that genetically separates the A and B lineages (Fig. 1). Based on association mapping in the chloroplast genome, the dual promoter region in the intergenic spacer between the *clpP* operon and the *psbB* operon was proposed to be involved in the incompatibility AB-I. The *clpP* gene encodes the proteolytic subunit of the Clp protease, the *psbB* operon encodes core subunits of photosystem II (PSII) and the cytochrome *b*_6_*f* complex (Cyt*b*_6_*f*)^31^. However, the molecular mechanism underlying the incompatibility have remained enigmatic.

## Results

### AB-I plants are unable to acclimate to higher light conditions

AB-I plants display a yellow-green *(lutescent)* leaf chlorosis, caused by disturbed PSII activity^31^ (Fig. 2a,b). The photosynthetic defects occur specifically under increased light intensities (Fig. 2b). Whereas at 300 μE m^-2^s^-1^ (low light, LL), the compatible wild type AB-II and the incompatible hybrid AB-I are indistinguishable from each other, higher light intensities cause severe photodamage in AB-I. Consistent with our previous study, already at 450 μE m^-2^s^-1^ (high light, HL), a substantial portion of PSII was photodamaged. Interestingly, AB-I plants are also unable to perform an efficient light acclimation response when shifted to HL conditions (Fig. 2b; Supplementary Text): Whereas AB-II plants responded to the increased growth light intensity by strongly increasing their chlorophyll content (Supplementary Figure 1a) and the contents of all redox-active components of the electron transport chain (Fig. 2b), AB-I plants were incapable of performing this light acclimation response efficiently. This behavior, inhibited in AB-I plants, is a typical reaction when plants previously grown under light-limited conditions are transferred to higher light intensities^32^. Finally, this leads to a relative reduction of the components of the electron transport chain, namely PSI, PSII and Cytb_6_*f*, but not the ATP synthase (ATPase) and plastocyanin (PC), in AB-I plants compared to AB-II (Fig. 2b,c; Supplementary Fig. 1, Supplementary Text). In summary, AB-I plants display a light-dependent phenotype of photosynthetic acclimation that cannot be assigned to a single component of the electron transport chain. In addition, the disturbance in acclimation response is independent of ATP synthase and PC function.

**Fig. 2.**
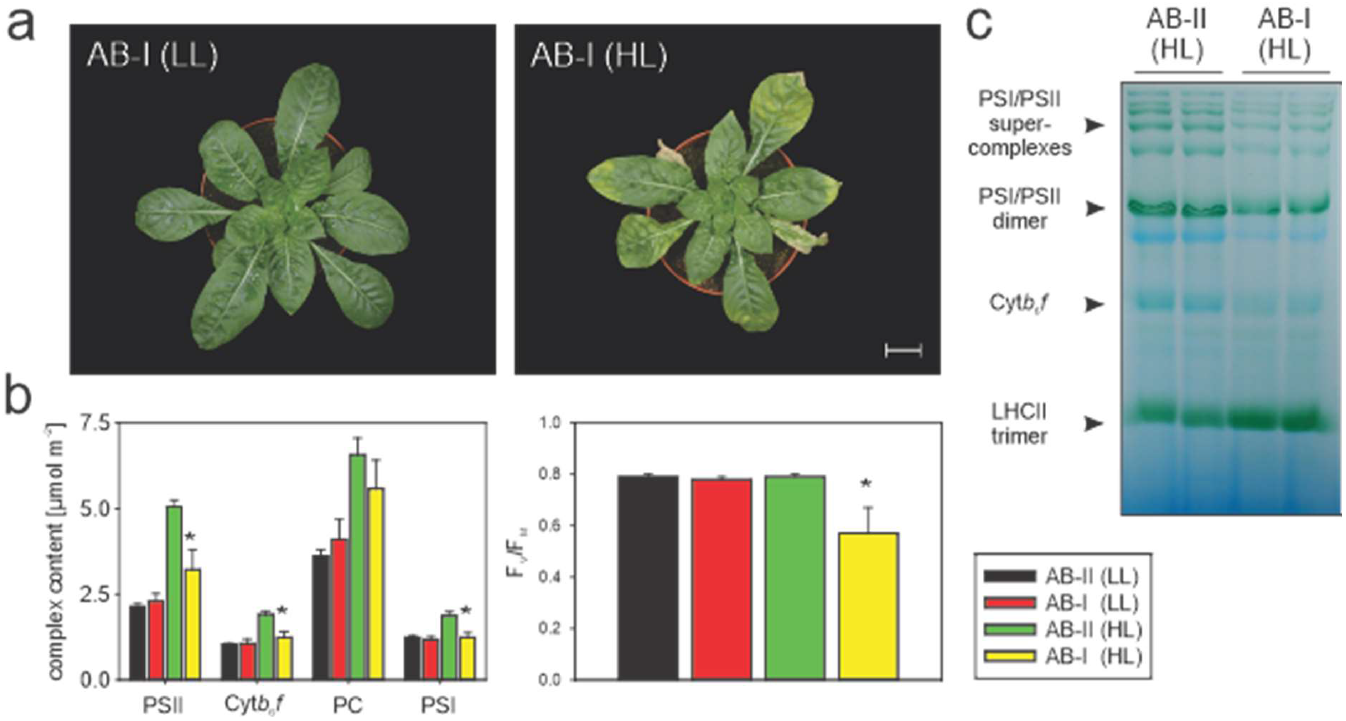
Light-dependent phenotype and physiology of AB-I plants. **a**, Yellow-green *(lutescent)* leaf phenotype and growth retardation under high-light (HL) condition (right panel). Scale bar: 5 cm. **b**, Left panel: Quantification of the components of the photosynthetic electron transport chain by difference absorbance spectroscopy. Note that AB-I plants under HL condition are not able to perform a typical light acclimation response by strongly increasing the contents of all redox-active components of the electron transport chain relative to low-light (LL) conditions. Bars represent mean values ±SD (n = 6-8). Asterisk indicates significant difference from AB-II HL t-test, *P* < 0.05 (PSII: t = 7.40, df = 12; Cyt*b*_6_*f*: t = 9.08, df = 12; PC: t = 2.59, df = 12; PSI: t = 8.88, df = 12). Right panel: Severe photooxidative damage of AB-I plants under HL conditions, exemplified by measurement of the maximum quantum efficiency of photosystem II in the dark-adapted state (F_V_/F_M_). Bars represent mean values ±SD (n = 6-8). Asterisk indicates significant difference from AB-II HL t-test, *P* < 0.05 (t = 5.15, df = 12). **c**, Bluenative PAGE independently confirming the reduction of the electron transport chain complexes in AB-I under HL. This experiment was performed independently two times with similar results.

### The better adaptation of wild-type AA-I plants to high light is conferred by the chloroplast genotype

To examine, if the genetic differences between plastome I and II have phenotypic effects under high light conditions also in a compatible background (i.e., could be subject to selection in the parental species), we compared the light-acclimation response of green wild-type AA-I (O. *elata)* plants with green wild-type AB-II (O. *biennis).* In addition, we investigated, whether potential differences are due to the chloroplast by including the green chloroplast substitution line of the two genotypes (AA-II; Fig. 1b). Since this experiment involved green material only, a harsher and more natural light shift from 300 μE m^-2^s^-1^ to 600 μE m^-2^s^-1^ (harsh high light, HHL) could be analyzed. The latter condition already induces severe damage in the incompatible AB-I genotype (Supplementary Text). In 300 μE m^-2^s^-1^, all photosynthetic parameters investigated (chlorophyll a/b ratio, chlorophyll content, F_V_/F_M_, linear electron transport capacity, and chloroplast ATP synthase activity) were very similar between the three genotypes (Table 1). However, after the shift to high light, pronounced differences started to occur. While most parameters were not, or only weakly, affected in AA-I plants, the AB-II genotype showed a drastic loss of electron transport capacity. This was accompanied by marked decreases in the chlorophyll a/b ratio, chlorophyll content per leaf area, and chloroplast ATP synthase activity. Strikingly, similar changes also occurred in AA-II plants, indicating that mainly the plastome, and not the nuclear genetic background, is causal for these differences in light acclimation. However, in comparison to typical light acclimation responses of angiosperms, which (due to degradation of the chlorophyll b binding antenna proteins) result in increased electron transport capacity, chlorophyll content and chlorophyll a/b ratio^32^, the response of AA-I plants to increased light intensity is limited. It, therefore, can be concluded that plastome I is better adapted to cope with high light conditions than plastome II, although at least under the conditions tested, the johansen Standard strain of *O. elata,* originally isolated in California^16,33^, does not behave like a typical high light plant.

**Table 1.** Comparison of light-acclimation responses of *O. elata* (AA-I), *O. bienns* (AB-II), and their green chloroplast substitution lines AA-II. Plants were either cultivated at 300 μE m^-2^ s^-1^ or 600 μE m^-2^ s^-1^ actinic light intensity, and their adaptive changes in chlorophyll a/b ratio, chlorophyll content per leaf areas, maximum quantum efficiency of PSII in the dark-adapted state (F_V_/F_M_), linear electron transport capacity (ETRII) and chloroplast ATP synthase activity (gH^+^) were compared between both light regimes by pair-wise testing.^1)^

| Parameter | AB-II 300$\mu\text{E}$: | AB-II 600$\mu\text{E}$: | AA-I 300$\mu\text{E}$: | AA-I 600$\mu\text{E}$: | AA-II 300$\mu\text{E}$: | AA-II 600$\mu\text{E}$: |
| --- | --- | --- | --- | --- | --- | --- |
| **Chlorophyll a/b** | 4,01 ($\pm\text{SD } 0,06$) | 3,60 $\pm\text{SD } 0,15^{**}$ | 4,08 $\pm\text{SD } 0,05$ | 4,04 $\pm\text{SD } 0,16$ | 3,92 $\pm\text{SD } 0,08$ | 3,74 $\pm\text{SD } 0,15^*$ |
| **Chl. [$\text{mg m}^{-2}$]** | 631,0 $\pm\text{SD } 18,1$ | 548,6 $\pm\text{SD } 104,3$ | 668,1 $\pm\text{SD } 27,1$ | 688,3 $\pm\text{SD } 58,8$ | 556,7 $\pm\text{SD } 8,4$ | 464,4 $\pm\text{SD } 34,1^{**}$ |
| **$F_v/F_m$** | 0,79 $\pm\text{SD } 0,01$ | 0,73 $\pm\text{SD } 0,08^*$ | 0,81 $\pm\text{SD } 0,01$ | 0,74 $\pm\text{SD } 0,06^*$ | 0,81 $\pm\text{SD } 0,01$ | 0,71 $\pm\text{SD } 0,03^{**}$ |
| **ETRII [$\mu\text{mol m}^{-2} \text{s}^{-1}$]:** | 172,3 $\pm\text{SD } 14,0$ | 75,1 $\pm\text{SD } 30,1^{**}$ | 171,5 $\pm\text{SD } 5,9$ | 137,5 $\pm\text{SD } 51,4$ | 147,9 $\pm\text{SD } 13,7$ | 84,4 $\pm\text{SD } 14,7^{**}$ |
| **$\text{gH}^+ [\text{s}^{-1}]$** | 39,2 $\pm\text{SD } 2,5$ | 27,4 $\pm\text{SD } 3,8^{**}$ | 43,0 $\pm\text{SD } 2,9$ | 45,8 $\pm\text{SD } 11,0$ | 40,4 $\pm\text{SD } 3,6$ | 33,3 $\pm\text{SD } 2,8^*$ |

<sup>1)</sup> Asterisks indicates significant difference ( $t$ -test, \* $P < 0.05$ , \*\* $P < 0.01$ , $n \geq 5$ )

### RNA editing is not involved in the AB-I incompatibility of *Oenothera*

Causative chloroplast loci for the described phenotypes could be related to mRNA editing sites that often display great variability between even closely related species^9,34^. RNA editing in chloroplasts of seed plants involves C-to-U conversions at highly specific sites^35^. It is of particular interest, since the only previously described mechanism of PGI is based on an editing deficiency of the tobacco *atpA* transcript (encoding a core subunit of the plastid ATP synthase) when exposed to the nuclear genetic background of the deadly nightshade *Atropa^9^.* However, as evidenced by sequencing of the chloroplast transcriptomes of *O. elata* (AA-I), *O. biennis* (AB-II), and *O. grandiflora* (BB-III), mRNA editing does not play a causal role in the AB-I incompatibility. All compatible wild-type genome combinations of the three species share the same 45 mRNA editing sites (Supplementary Table 2 and updated GenBank records AJ271079.4, EU262889.2, and KX014625.1). This analysis also includes partially edited sites, whose biological relevance is doubtful (see Supplementary Table 2 and Materials and Methods for details). These results exclude the possibility that editing sites in the plastome and/or nucleus-encoded editing factors differ between the genotypes involved in the AB-I incompatibility, as this was reported for an experimentally produced cybrid of *Atropa* and tobacco^9^.

### Association mapping of plastid loci causing the AB-I incompatibility

Having ruled out the involvement of mRNA editing, we performed an association mapping in the chloroplast genome of *Oenothera* to pinpoint the causative loci for the AB-I incompatibility. In contrast to the green alga *Chlamydomonas,* chloroplast genomes of higher plants are not amenable to linkage mapping^36,37^. Hence, identification of functionally relevant loci is usually based on correlation of a polymorphism to a phenotype in a mapping panel (e.g. Refs.^31,38,39^). In the case of the AB-I incompatibility, this can be achieved by manual inspection of an alignment of fully sequenced chloroplast genomes and search for specific polymorphisms in plastome I vs. II, III and IV. Those polymorphisms are considered candidates for causing the AB-I incompatibility, because only plastome I confers the bleached *lutescent* phenotype in the AB nuclear genetic background, whereas plastomes II, III and IV are all green when combined with the same nucleus^15,31^. Our original analyses of the AB-I phenotype had included only four chloroplast genomes and yielded 16 candidate regions^31^. Taking advantage of the power of next-generation sequencing technologies, we now were able to base the association mapping on 46 full chloroplast genomes, whose genetic behavior had been determined by extensive crossing studies^40–43^ (Methods; Supplementary Table 1). The chosen strains represent the material used for generalization of the genetic species concept in *Oenothera* that is based on the basic A, B and C nuclear and I – V chloroplast genotypes^16,21^ (see Introduction). Altogether, the mapping panel included 18 chloroplast genomes representing plastome type I (Methods; Supplementary Table 1). Only four polymorphisms were absolutely linked with the AB-I phenotype, in that they were specific to plastome I and could potentially be involved in the AB-I incompatibility: (i) a 144 bp deletion in the *clpP* – *psbB* operon spacer region, (ii) a combined 5 bp deletion/21 bp insertion (indel) in the *psbM* – *petN* spacer (genes encoding a PSII and a Cytb_6_*f* subunit, respectively), (iii) a 194 bp deletion in the *ndhG* – *ndhI* spacer (two genes encoding subunits of NADH dehydrogenase complex), and (iv) a 21 bp insertion in the *trnL-UAA* – *trnT-UGU* spacer (Supplementary Datasets 1-3).

Due to the lack of measurable sexual recombination frequencies in chloroplast genomes of seed plants^37^ (see above), genetic methods cannot be employed to further narrow down on the causative loci for the AB-I incompatibility in plastome I. We, therefore, evaluated the remaining candidate polymorphisms with respect to their potential to cause the incompatible phenotype. The deletion in the *ndhG* – *ndhI* spacer and the deletion in the *trnL-UAA* – *trnT-UGU* spacer cannot explain the observed light-dependent reduction of specific photosynthetic complexes in AB-I incompatible material (Fig. 2). The neighboring genes do not encode components of the electron transport chain, and, moreover, knockouts of NDH complex subunits lack any discernible phenotype^44^. Possible effects on the expression of *trnL-UAA* and/or *trnT-UGU,* two essential tRNAs, would be much more pleiotropic and not depend on the light intensity. Based on the functions of the genes affected, a contribution of the latter two polymorphisms to the AB-I phenotype is extremely unlikely. By contrast, the polymorphisms affecting the *psbB* operon and the *psbM*/*petN* spacer are serious candidates, in that they potentially affect both PSII and Cytb_6_*f*, which is in line with the physiological data (Fig. 1b).

### The *psbN-petN* spacer region may make a minor contribution to the AB-I incompatibly

To examine the contribution of the combined 5 bp/21 bp indel in the *psbM* – *petN* spacer, transcript and protein analyses were performed in incompatible AB-I plants and compatible controls under LL and HL conditions (Fig. 3). The indel is located in the 3’-UTR of both genes (Fig. 3a) and, therefore, could potentially affect the stability of their transcripts.

**Fig. 3.**
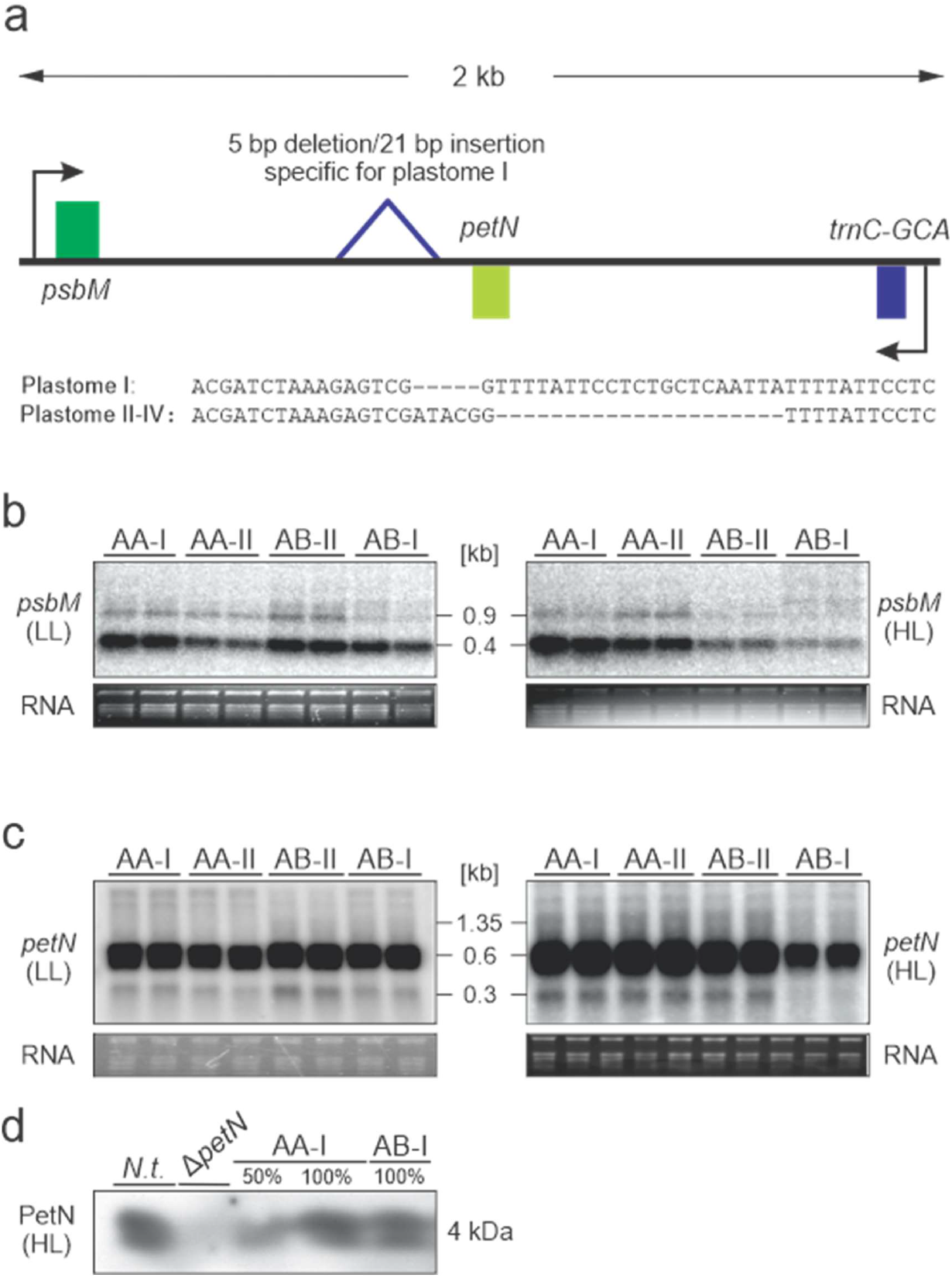
Molecular genetic analyses of the *psbM* – *petN* spacer region in compatible (AA-I, AA-II, AB-II) and incompatible material (AB-I) under HL and LL conditions. **a**, Sequence context and insertions/deletions in the spacer that are specific to plastome I. Arrows indicate transcription start sites. Northern blot analysis of *psbM* (**b**) and *petN* (**c**) transcript accumulation under LL and HL conditions. These experiments were performed independently three times with similar results. **d**, Western blot analysis of PetN protein accumulation under HL. *N.t.* = *Nicotiana tabacum* WT, *ΔpetN* = *petN* knockout in *Nicotiana tabacum^47^.* The experiment was performed independently three times with similar results.

Northern blot analyses revealed that both genes are affected by the indel. For *psbM,* reduction of the 0.35 kb monocistronic transcript was observed for AA-II and AB-I under LL, and for AB-II and AB-I under HL conditions. Although there is no obvious explanation for this light-dependent effect, it is independent of the AB-I incompatibility in that the levels of *psbM* mRNA cannot be linked to the AB-I phenotype (Fig. 3b and below). Moreover, as judged from knockout mutants in tobacco, even complete loss of the PsbM protein does not lead to a strong phenotype that would be comparable to our material^45^. By contrast, *petN* encodes an essential subunit of the Cytb_6_*f*^46,47^, and reduced *petN* transcript stability, therefore, could affect Cytb_6_*f* accumulation. Northern blot analysis of *petN* mRNA accumulation detected a mature transcript of 0.3 kb (Fig. 3a). Under LL conditions, *petN* transcript accumulation is unaltered in the incompatible hybrid, whereas under HL, the *petN* mRNA is significantly reduced in AB-I material (Fig. 3c). Western blot analyses showed that this leads to a reduction at the protein level to approximately 80% (Fig. 3d), an estimate that is well supported by our spectroscopic quantification of Cytb_6_*f* (Fig. 1b).

Taken together, these data do not exclude the possibility that the *psbM/petN* region influences the incompatibility phenotype, but suggest a rather minor contribution. First, involvement of *psbM* is very unlikely, because down-regulation of its mature transcript is observed also in compatible AA-II plants under LL conditions (Fig. 3b). Second, a role of *petN* is unlikely as well, since reduction of about 20% of the Cyt*b*_6_*f* content (Fig. 2b, Fig. 3d) does not affect accumulation of the photosystems^46–49^. Hence, another chloroplast locus must be causally responsible for the AB-I incompatibility.

### The promotor region of the *psbB* operon is the major locus causing the AB-I incompatibly

Next, we analyzed the transcript patterns of the *clpP* and *psbB* operons that flank the 144 bp deletion in the spacer region (Supplementary Fig. 2a). Northern blot analyses revealed that accumulation of both the *clpP* precursor transcript and the mature *clpP* mRNA did not differ in control plants and incompatible plants under HL conditions. All *clpP* transcripts accumulated to similar levels as in the compatible lines. Similarly, no difference in transcript accumulation of the remaining operon genes residing upstream of *clpP (rpl20* and *5’-rps12)* was observed (Supplementary Fig. 2b). In addition, analyses of ClpP protein accumulation and the integritiy of the plastid ribosomes revealed no difference between compatible and incompatible material^50^. Based on these findings, a contribution of the *clpP* operon to the incompatibility phenotype can be excluded. By contrast, transcript accumulation of all *psbB* operon genes *(psbB, psbT, psbH, petB* and *petD)* was found to be reduced in AB-I plants under HL, but not under LL conditions (Fig. 4b,f; Supplementary Fig. 2). Run-on transcription analyses revealed that this effect was due to impaired transcription rather than an effect of altered transcript stability. *psbB* operon transcription was specifically reduced under HL conditions in the incompatible hybrids (Fig. 4d). Consequently, in contrast to the green AA-I, AA-II and AB-II plants, the deletion in plastome I in the AB background affects regulation of the *psbB* operon promoter in a light-dependent manner (Fig. 4d). Importantly, the same promoter is used in all genetic backgrounds, as evidenced by mapping of the transcription start sites (Fig. 4c), which are also highly conserved between species (Fig. 4a). The deletion does not affect the TATA box of the *psbB* operon promoter, but resides 7 bp upstream of the −35 box. This may suggest that polymerase binding *per se* is not affected, and instead, binding of auxiliary proteins such as sigma factors is impaired by the deletion in the incompatible hybrids (see Discussion).

**Fig. 4.**
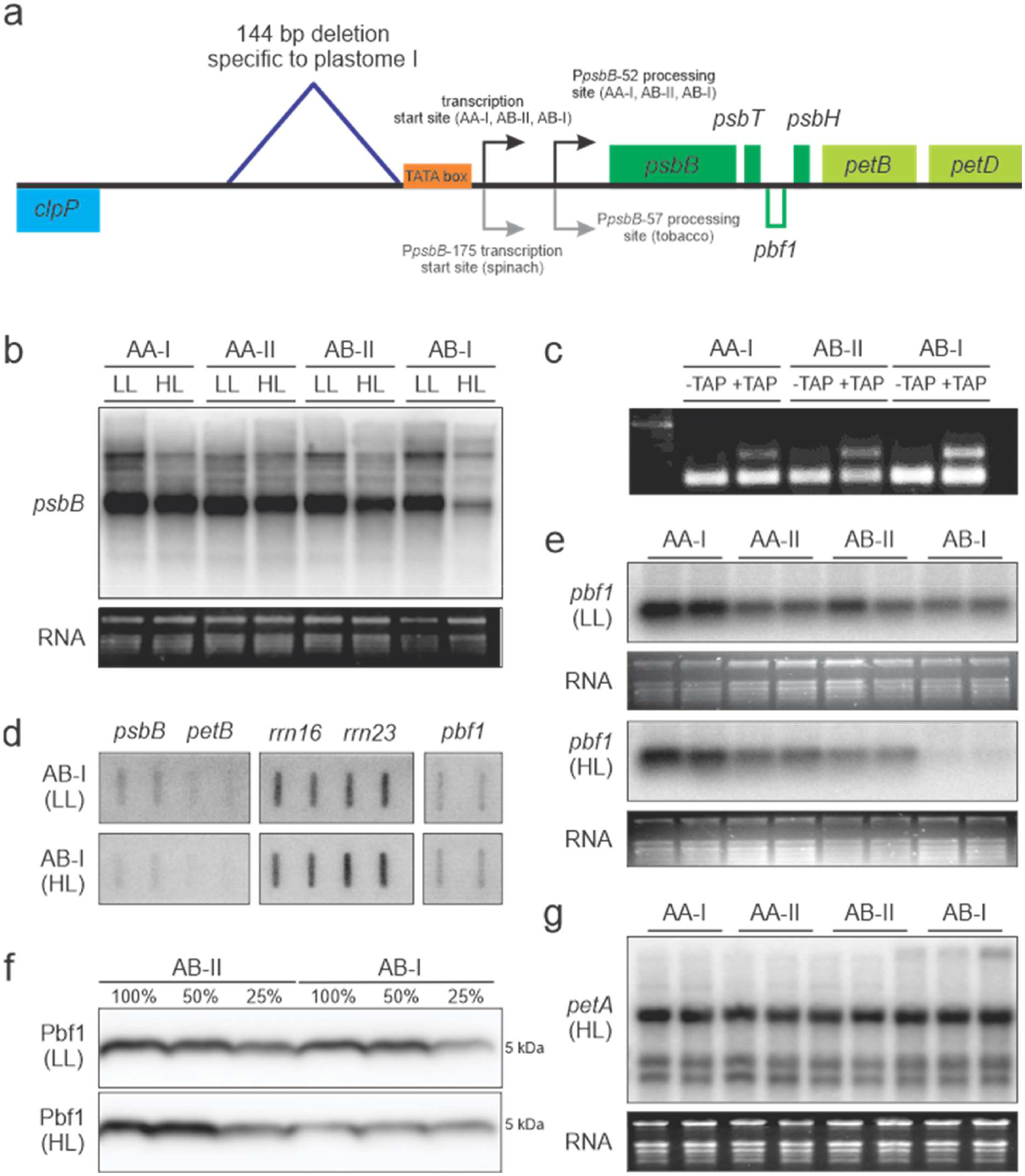
Regulation of the *psbB* operon in compatible (AA-I, AA-II, AB-II) and incompatible plants (AB-I) under HL and LL conditions. **a,** Physical map of the region in the chloroplast genome containing the *clpP* and *psbB* operons. The 148 bp deletion in the intergenic spacer upstream of the *psbB* operon promoter is indicated. Transcription start sites and mRNA processing sites are indicated. Note that the *pbf1* gene (encoded on the opposite strand) is transcribed from its own promoter. **b,** Northern blot analysis of *psbB* transcript (representative of the whole *psbB* operon; also see Supplementary Fig. 2). The experiment was performed independently three times with similar results. **c,** 5’-RACE with and without TAP treatment, a method to map transcription start sites and RNA processing sites of the *psbB* operon. For details see Methods. This experiment was performed independently three times with similar results. **d,** Run-on transcription analysis of the *psbB* operon *(psbB, petB),* appropriate controls *(rrn16, rrn23),* and *pbf1.* The experiment was performed independently three times with similar results. **e,** Northern blot analysis of *pbf1* transcripts. The experiment was performed independently three times with similar results. **f**, Western blot analysis of Pbf1 protein accumulation. The experiment was performed independently two times with similar results. **g,** Northern blot analysis of *petA* transcript accumulation, serving as a control for a gene outside of the *psbB* operon. The experiment was performed independently three times with similar results.

Interestingly, *pbf1 (photosystem biogenesis factor 1,* previously designated *psbN),* a gene involved in PSI and PSII assembly, is down-regulated in AB-I incompatible plants under HL conditions (Fig. 4e). Since the gene is transcribed from the opposite strand by its own promoter that lacks any polymorphism in all *Oenothera* plastomes sequenced so far (Supplementary Datasets S1-S3), the reduction in *pbf1* transcript accumulation must result from the sense-antisense interaction with the *psbT* mRNA, as previously described for *Arabidopsis^51,52^.* Alternatively, it might be the result of an unknown feedback regulation. In any case, the interaction results in a strong reduction of Pbf1 protein accumulation (Fig. 4f). Since *pbf1* knockouts are extremely light-sensitive, and show severe defects in PSII and, to a lesser extent, also in PSI accumulation^53,54^, it appears likely that the effect on the *pbf1* mRNA also contributes to the incompatibility phenotype.

## Discussion

Our work reported here shows that light-dependent misregulation of a core photosynthesis operon leads to hybrid incompatibility, thus causing reproductive isolation and ultimately, speciation. Interestingly, the underlying genetic architecture was shaped during the last ice age by periods of glaciation (see Introduction). The mechanism we have uncovered is different from that of the two other PGIs studied so far. RNA editing of the *atpA* transcript was identified as causal for chloroplast-nuclear incompatibly in an *Atropa*/tobacco synthetic cybrid^9^ (see above). Variation in the coding regions of *accD* (the plastid-encoded subunit of the acetyl-CoA carboxylase, catalyzing the first step of fatty acid biosynthesis) was suggested as a genetic determinant of PGI in pea^55^. However, the Atropa/tobacco case represents a non-natural, artificial combination of the plastid and the nuclear genomes of two non-crossable species, and, unfortunately, the evidence for possible causative loci for the incompatibility in pea are currently not strong enough to judge their impact on natural populations^56^. Moreover, in both cases, the ecological relevance of the suggested PGI loci is unclear and cannot be deduced from the identified polymorphisms.

By contrast, our data indicate that the AB-I incompatibility might have evolved as a result of ecological selection. That cytoplasmic incompatibly can result from ecological selection is obvious from work in sunflower, were common garden experiments in xeric and mesic habitats demonstrated maintenance of cytoplasmic incompatibility by positive selection^57^. The underlying genes and physiology, however, have remained enigmatic. Our study demonstrates that photosynthesis-related genes encoded in the chloroplast genome can establish hybridization barriers. The incompatible phenotype is only visible under HL condition in that AB-I plants cannot perform the necessary acclimation response (see above). Strikingly, plastome I in the native AA nuclear genetic background of *O. elata* (a species adapted to the western United States and Mexico) copes better with high light conditions than the AB-II genotype of *O. biennis,* native to the North American woodlands^16^. This effect is plastome dependent, since the green AA-II chloroplast substitution line is phenotypically very similar to *O. biennis.* However, to what extent the identified incompatibility locus is involved in a light acclimation response of AA-I species in their natural habitats remains to be addressed in further investigations. At least within their natural range of distribution, species carrying plastome I colonized Central and the south west of North American (xeric habitats with high light irradiation), whereas species carrying plastomes II, III, IV or V are limited to the mesic sites of eastern North America (Fig. 1C)^16,21^. Hence, plastome I seems to be a required for colonization of habitats exposed to higher irradiation. This assumption is further supported by the fact that *O. biennis* (AB-II or BA-III), a species that spread after 1970 west of the Great Plains, is only rarely found in the southern parts of the USA and is still absent from Mexico^16,21^. Consequently, the loci underlying the AB-I incompatibility seem to prevent colonization of the south western parts of North America by the B genome by creating an asymmetric hybridization barrier between AA-I and AB-II, BA-III and BB-III species. At the same time, the compatible genome combination AA-I may have facilitated physiological adaptation of the corresponding species by nuclear-cytoplasmic co-evolution. It, therefore, seems reasonable to assume that, as a result of higher light intensities (and/or light quality differences) in xeric habitats, the deletion upstream of the *psbB* operon promoter co-evolved with nucleus-encoded proteins that interact with the (bacterial-type) plastid-encoded RNA polymerase (PEP).

Strong candidates for these interacting proteins are the PEP sigma factors, which were shown to regulate polymerase binding in response to both light quality and light quantity^58,59^. Moreover, regulation by sigma factors (e.g., through redox induced phosphorylation) influences the stoichiometry of the protein complexes of the photosynthetic electron transport chain^60^. Thus, the failure of AB-I plants to acclimate to high light intensities could be a direct consequence of disturbed transcriptional regulation by sigma factors. Co-evolution and coordinated rates of molecular evolution of PEP core subunits and sigma factors could be a common principle in plant evolution, as suggested by recent findings in Geraniaceae, a family where PGI is also widespread^61^.

Finally, it should be emphasized that, although the *psbB* operon does not encode PSI-related genes, its transcriptional misregulation also explains the observed effect on PSI, due to the antisense interaction with the *pbf1* mRNA or an unknown mechanism of feedback regulation (Fig. 4). Interestingly, while the *psbB* operon (including the *pbf1* gene on the opposite strand) displays extremely high structural conservation from cyanobacteria to higher plants, light regulation of *pbf1* transcript abundance was shown to be highly variable between species^62^.

## Methods

### Plant material

Throughout this work, the terms *“Oenothera”* or “evening primrose” refer to subsection *Oenothera* (genus *Oenothera* section *Oenothera,* Onagraceae; 2n = 2x = 14)^16^. Plant material used here is derived from the *Oenothera* germplasm resource harbored at the Max Planck Institute of Molecular Plant Physiology (Potsdam-Golm, Germany), which includes the living taxonomic reference collection of subsection *Oenothera^17^.* Part of this reference collection is the so-called Renner Assortment, a medium-sized collection of European lines thoroughly characterized by the genetic school of Otto Renner^21,63^. In addition, it includes the Cleland collection, a large set of North American strains of subsection *Oenothera* that was extensively studied by Ralph E. Cleland^21^. Present as well are North American accessions analyzed by Wilfried Stubbe and co-workers, which represent the species of this subsection recognized later than the 1960s^64–71^. The availability of this material allowed us to employ the original source of lines on which the genetic species concept of subsection *Oenothera* is based cf. ref.^72^ (see Main Text). The lines employed for association mapping of the plastidic AB-I locus were extensively analyzed by classical genetics for the compatibility relations of their nuclear and chloroplast genomes (Supplementary Table 3 for details).

RNA editing analyses were performed with the wild type strains of *Oenothera elata* subsp. *hookeri* strain johansen Standard (AA-I), *O. grandiflora* strain Tuscaloosa (BB-III) and *O. biennis* strain suaveolens Grado (AB-II). See Supplementary Table 3 for a summary of all wild-type strains used in this work.

For all other genetic or physiological work presented here, the wild type strains johansen Standard (AA-I) and suaveleons Grado (AB-II) or chloroplast substitution lines between them (AA-II and AB-I) were used. Here, AA-I refers to the wild type situation, i.e. strain johansen Standard with its native nuclear and chloroplast genomes. AA-II refers to the nuclear genome of johansen Standard combined with the chloroplast genome of suaveolens Grado. AB-II designates nuclear and chloroplast genomes of the wild type strain suaveolens Grado, and AB-I the nuclear genome of suaveolens Grado equipped with the chloroplast genome of johansen Standard. Generation of AA-II and AB-I from the wild types AB-II and AA-I is detailed below (also see summary in Supplementary Table 4).

As tobacco wild type, the cultivar Petit Havana was used. The tobacco *ΔpetN* mutant was obtained from^47^.

### Generation of chloroplast substitution lines

In *Oenothera,* the genetics of permanent translocation heterozygosity, combined with a biparental transmission of plastids offer an elegant opportunity to substitute chloroplasts between species in only two generations, while leaving the nuclear genome constitution unaltered^40,41,73^. The general principles, including a detailed discussion of crossing examples, are presented for example in Rauwolf *et al.* (2008)^22^. The interested reader is referred to this previous work. The chloroplast substitution between the strains suaveolens Grado and grandiflora Tuscaloosa [described in Fig. 6 of Rauwolf *et al.* (2008)^22^] resembles the chloroplast substitution between suaveolens Grado and johansen Standard used in this work.

In brief, due to reciprocal chromosomal translocations, many species of *Oenothera* form permanent multi-chromosomal meiotic rings. If all members of a given chromosome complement are involved in a single ring, they establish two regularly segregating sets of genetically linked chromosomes. This leads to formation of two superlinkage groups, each involving one complete parental haploid chromosome set (α and β). Suppression of homologous recombination avoids genetic reshuffling between the two haploid sets. Additional genetic properties, especially presence of gametophytic lethal factors that lead to sex-linked inheritance of a given haploid set, eliminate homozygous segregants (α·α or β·β). This results in permanent heterozygous progeny (α·β) that is identical to the parental plant. The phenomenon of structural heterozygosity is a form of functional “asexuality”. However, also “sexual” species exist in *Oenothera,* i.e., species that display bivalent-pairing and regular meiotic segregation. In contrast to the structurally heterozygous species, they lack lethal factors and are homozygotous for their haploid sets (haplo·haplo vs. α·β from above)^21,63^.

As a consequence of this genetic behavior, entire haploid chromosome sets in evening primrose can behave as alleles of a single Mendelian locus. These so-called Renner complexes are designated with (Latin) names; e.g. *^G^albicans·^G^flavens* (α·β) for the structurally heterozygous strain suaveolens Grado or *^h^johansen Standard. ^h^johansen Standard* (haplo·haplo) for the homozygous line johansen Standard. A cross between them (suaveolens Grado x johansen Stanard = *^G^ablicans. ^G^flavens* x *^h^johansen Standard. ^h^johansen Standard*) yields in the F1 the offspring *^G^albicans. ^h^johansen Standard* and *^G^flavens. ^h^johansen Standard*.

To equip johansen Standard (AA-I) with the chloroplast of suaveolens Grado (AB-II), the F1 hybrid *^G^albicans.^h^johansen Standard* is used. It carries (due to biparental inheritance of chloroplasts in evening primroses) the chloroplasts of both johansen Standard (I-johSt) and suaveolens Grado (II-suavG). (The other hybrid *^G^flavens.^h^johansen Standard* is not of interest and therefore discarded.) Since *^G^albicans.^h^johansen Standard* I-johSt/II-suavG displays a full meiotic ring, this leads to suppression of homologous recombination as well as elimination of random chromosome assortment in meiosis (see above). Therefore, as a result of Mendelian segregation of the *^h^johansen Standard* complex, the johansen Standard strain (*^h^johansen Standard.^h^johansen Standard*) can be bred back from *^G^albicans.^h^johansen Standard* upon selfing. (*^G^albicans.^h^johansen Standard* x s = *^G^albicans.^G^albicans, ^G^albicans.^h^johansen Standard* and *^h^johansen Standard.^h^johansen Standard*; the sergeant *^G^albicans.^G^albicans* is not realized due to a male gametophytic lethal factor in *^G^albicans*). When a *^G^albicans.^h^johansen Standard* plant homoplasmic for II-suavG is used for selfing, the johansen

Standard plant in F2 now carries plastome II-suavG (AA-II). If AB-I plants are desired, the *^G^albicans.^h^johansen Standard* I-johSt/II-suavG hybrid in F1 is selected for I-johSt and backcrossed with suaveolens Grado (*^G^abicans.^G^flavens* II-suavG). BC1 then reassembles *^G^albicans.^G^flavens* I-johSt/II-suavG which, due to the maternal dominance of biparental transmission in evening primrose^39,74,75^, contains a major proportion of *^G^albicans.^G^flavens* I-johSt, i.e., AB-I plants.

### Plant cultivation, growth conditions, and tissue harvest

For crossing studies, plastome sequencing and analysis of RNA editing, *Oenothera* plants were cultivated in a glasshouse as previously described^17^. AA-I, AA-II, AB-I and AB-II plants for genetic and physiological analyses were cultivated in soil in growth chambers at a 16 h light/8 h darkness cycle and 24°C at low light intensities (~150 μE m^-2^s^-1^). After formation of the early rosette (cf. ref.^17^), plants were transferred to higher light intensities, i.e., 300 μE m^-2^s^-1^ (low light, LL), 450 μE m^-2^s^-1^ (high light, HL) or 600 μE m^-2^s^-1^ (harsh high light, HHL), and kept under the same growth regime. 600 μE m^-2^s^-1^ was used only for a single experiment, because it already resulted in severe photodamage of the incompatible combination AB-I (see Supplementary Text). To avoid pleiotropic effects, the yellowish material of the bleached leaf tip, a typical characteristic of the *lutescent* AB-I incompatible phenotype (Fig. 1a), was excluded from all experiments. The tobacco *ΔpetN* mutant and its corresponding wild type were cultivated as reported earlier^47^.

### Thylakoid membrane isolation from *Oenothera* leaves

For spectroscopic measurements and bluenative PAGE, an improved thylakoid membrane isolation protocol was developed for *Oenothera* leaf tissue that contains high amounts of mucilage and starch. All steps were performed at 4°C. Solutions were pre-chilled, leaves shortly placed in ice-cold water and dried with a salad spinner. Approximately 10 g of mature leaf tissue dark adapted for 1 h was homogenized in a blender adding 200 ml of Isolation Buffer [330 mM sorbitol, 50 mM HEPES, 25 mM boric acid, 10 mM EGTA, 1 mM MgCl_2_, 10 mM NaF (optional), pH 7.6 with KOH, and 5 mM freshly added Na-ascorbate]. 100 ml aliquots of the homogenate were then filtered through a double layer of cheese cloth (Hartmann), followed by filtering through a single layer of Miracloth (Merck). After that, the following procedure was applied twice: After adjustment of the solution to 200 ml with Isolation Buffer, it was centrifuged for 5 min at 5,000 g and the pellet subsequently resuspended in 40 ml of Isolation Buffer using a 30-cm^3^ Potter homogenizer (mill chamber tolerance: 0.15 to 0.25 mm; VWR). Subsequent to the second homogenization step, the solution was adjusted to 200 ml with Washing Buffer (50 mM HEPES/KOH pH 7.6, 5 mM sorbitol, and optionally 10 mM NaF) followed by a filtering step through one layer of Miracloth. Subsequently, the thylakoid homogenate was centrifuged for 5 min at 5,000 g, the pellet resuspended with a 30-cm^3^ Potter homogenizer in 30 ml Washing Buffer and centrifuged for 5 min at 5,000 g. Then, after resuspending the thylakoids in 5 ml of Washing Buffer, the homogenate was placed on a 85% Percoll cushion [Percoll stock solution: 3% (w/v) polyethylene glycol 6000, 1% (w/v) BSA, 1% (w/v) Ficoll 400, dissolved in Percoll; 85% Percoll: 85% PBF-Percoll stock solution, 330 mM sorbitol, 50 mM HEPES, 2 mM EDTA, 1 mM MgCl2; pH7.6 with KOH] in a 30 ml Corex tube and centrifuged for 5 min at 5,000 g. This step effectively removes starch from the isolation. Finally, thylakoids (that do not enter the Percoll cushion) are collected, washed in altogether 25 ml of Washing Buffer, centrifuged for 5 min at 5,000 g and resuspended in the desired puffer and volume.

### Spectroscopic methods

For quantification of isolated thylakoids, chlorophyll amounts were determined in 80% (v/v) acetone^76^. The contents of PSII, PSI, Cyt*b*_6_*f* and PC were determined in thylakoids as described previously^77^. PSI was quantified from P700 difference absorption signals at 830 to 870 nm in solubilized thylakoids using the Dual-PAM-100 instrument (Walz)^49,78^. Contents of PSII and Cyt*b*_6_*f* were determined from difference absorption measurements of cytochrome b559 and Cyt*b*_6_*f*, respectively. Measurement procedures and data deconvolution methods have been described previously in detail^78,79^. Maximum F_v_/F_m_ values were measured in leaves adapted for one hour to darkness. Chlorophyll fluorescence was recorded with a pulse amplitude-modulated fluorimeter (Dual-PAM-100) on intact plants at room temperature. A F-6500 fluorometer (Jasco) was used to measure 77 K chlorophyll-a fluorescence emission spectra on freshly isolated thylakoid membranes equivalent to 10 μg chlorophyll ml^-1^. The sample was excited at 430 nm wavelength with a bandwidth of 10 nm, and the emission spectrum was recorded between 655 and 800 nm wavelengths in 0.5 nm intervals with a bandwidth of 1 nm. Dark-interval relaxation kinetics of the electrochromic shift, which is a measure for the proton motive force across the thylakoid membrane, were used to determine the thylakoid conductivity for protons (gH^+^), which is a proxy for ATP synthase activity. Electrochromic shift signals were measured and deconvoluted using a KLAS-100 spectrophotometer (Walz) as previously described^80^.

### Antibody source and anti-PetN serum production

The Pbf1 (PsbN) antibody used in this work was described in Torabi *et al.* (2014)^54^. The anti-AtpA antibody and the secondary antibody (anti-rabbit IgG peroxidase conjugate antibodies) were obtained from Agrisera. To prepare an antibody against the PetN protein, rabbits were injected with PEG2-FTFSLSLVVWGRSGL-PEG2-C-Amid (BioGenes GmbH), a highly hydrophobic peptide comprising about half of the PetN protein. The peptide was coated with PEG2 (8-amino-3,6-dioxaoctanoic acid) to ensure better solubility. Active serum was obtained after four immunizations.

### Protein analyses

Blue-native PAGE was performed as previously reported^81,82^. To avoid protein degradation, 10 mM of NaF was added to all solutions. Thylakoid membranes were solubilized with dodecyl-β-D-maltoside (DDM) at a final concentration of 1% and separated in 4-12% polyacrylamide gradient gels. Protein equivalents of 30 μg chlorophyll were loaded.

For western blot analyses, thylakoids were mixed with Sample Buffer [50 mM Tris/HCl, pH 6.8, 30% (v/v) glycerol, 100 mM DTT, 4% (w/v) SDS, 10% (w/v) Coomassie Brilliant Blue G-250] and denatured for 5 min at 95°C under continuous agitation. Then, samples were subjected to Tricine-SDS-PAGE (16% T separation gel and 4% T stacking gel) followed by gel blotting onto a PVDF membrane (0.2 μm) using the semi-dry PEQLAB transfer system (PEQLAB Biotechnologie GmbH). After incubation with the secondary antibody, immunochemical detection was performed with the help of the ECL Prime Western Blotting Detection Reagent (GE Healthcare) according to the supplier’s recommendations. In the relevant figures, 100% loading corresponds to 3 μg chlorophyll equivalent.

### Isolation of nucleic acids

DNA and RNA isolations from evening primroses were performed employing protocols specially developed for their mucilage and phenolic compound rich tissue, as previously described in Massouh et al. (2016)^83^.

### Association mapping of the plastid AB-I locus

For association mapping in the chloroplast genome, 46 full plastome sequences of *Oenothera* were employed for which precise genetic information is available (Supplementary Table 1 and Plant Material section). To this end, we newly determined the sequences of 30 plastomes, now available from GenBank under the accession numbers KT881175.1, KX014625.1, MN807266.1, MN807267.1, and MN812468.1 to MN812493.1. The new chloroplast genomes were annotated and submitted by GeSeq v1.43 and GB2sequin v1.3^84^, respectively. The remaining 16 plastomes were previously published (see Supplementary Table 1 for details). Chloroplast genome sequencing from *Oenothera* total DNA was done as reported earlier^39,83^, but using a higher version of the SeqMan NGen assembly software (v14.1.0; DNASTAR). Also, in contrast to earlier work, 250 bp Illumina paired-end reads (instead of 100 bp or 150 bp) were generated, with the exception of KX014625.1 (100 bp paired-end) and MN807266.1 and MN807267.1 (both 150 bp paired-end). Subsequently, for association mapping, the redundant inverted repeat A (IR_A_) was removed, sequences were aligned with ClustralW and the alignments manually curated in Mesquite v3.40^85^. Polymorphisms specific to plastome I (i.e., polymorphisms that were present in all 18 plastome I genotypes, but absent from all 28 plastome II, III and IV genotypes) were identified by visual inspection in SeqMan Pro v15.2.0 (DNASTAR) cf. ref.^31^. For original data, see Supplementary Dataset 1.

### RNA editing analyses

To determine the RNA editotype of the *Oenothera* chloroplast, RNA-seq samples of the 1kp project^28,86^ of johansen Standard (AA-I), suaveolens Grado (AB-II), and grandiflora Tuscaloosa (BB-III; NCBI SRA Accession Numbers ERS631151, ERS631122, and ERS631139; also see Plant Material section) were mapped against their respective chloroplast genomes (AJ271079.4, KX014625.1, and EU262889.2) from which the IR_A_ had been removed. For this we employed the “reference-guided assembly – special workflows” pipeline of SeqMan NGen v15.2.0. SNPs were called in SeqMan Pro v15.2.0. To deal with the heterogeneity of the mRNA population, partial editing and sequencing errors, sites showing C-to-T (U) conversion of at least 30% were originally considered as mRNA editing sites. If editing could not be detected above this threshold at a given site in all three species, the sites were subjected to manual inspection of the original mapping data. In most cases, this procedure revealed mapping errors, however, in a few cases also partial editing below 30% in at least one of the strains was uncovered.

### Gel blot detection of RNA

Northern blot analyses were performed as previously described^83^. Genespecific PCR products used as probes were obtained by employing the primers listed in Supplementary Table 5. Total *Oenothera* DNA was used as template in standard PCR reactions.

### Chloroplast run-on analyses

For slot-blot preparation of DNA probes, PCR-amplified DNA probes (Supplementary Table 5) were immobilized to a Hybond-N+ nylon membrane (Amersham) through a slot-blot manifold. For this, 1.5 μg of DNA was denatured in 0.5 M NaOH and heated for 10 min at 95°C. Then, the volume of the denatured DNA probes was adjusted with water to 100 μl per spot. After heating, the probes were cooled on ice for 2 min to prevent DNA renaturation, and briefly centrifuged to collect the condensate. To each sample, 20 μl of cold 0.5 M NaOH and 0.5 μl of cold 10 x DNA-Loading Dye [50% (v/v) glycerol, 100 mM EDTA, 0.25% (w/v) bromophenol blue, 0.25% (w/v) xylene cyanol] were added. Subsequently, the samples were spotted to the ddH2O pre-hydrated nylon membrane and then 100 μl 0.5 M NaOH was applied to each spot. After drying the membrane at room temperature for 5 min, the DNA was cross-linked to the membrane with 0.12 J/cm^2^ using the UV crosslinker BLX-254 (BIO-LINK).

To analyze strand-specific gene expression of *pbf1,* single-stranded *pbf1* RNA probes were generated using the Ambion^®^ Maxiscript^®^ T7 Kit (Invitrogen) according to the manufacturer’s instructions and immobilized through a slot-blot manifold to a Hybond-N nylon membrane. The *pbf1* gene of johansen Standard was amplified with the primer pair psbNRO_F 5’-AGCATTGGGAGGCTCATTAC-3’ and psbNRO_R 5’-GGAAACAGCAACCCTAGTCG-3’ and cloned into to pCRTM2.1-TOPO^®^ (Invitrogen). The vector was linearized with *Hind*III and *in vitro* transcription was performed according to the suppliers’ protocol. 1.5 μg of RNA was adjusted with nuclease-free water to a volume of 50 μl prior to incubation with 30 μl of 20 x SSC (0.3 M sodium citrate and 3.0 M sodium chloride) and 20 μl 37% formaldehyde at 60°C for 30 min. Samples were maintained on ice and spotted to the ddH2O and 10 x SSC pre-hydrated nylon membrane. Next, 100 μl 10x SSC was applied per slot. After drying the membrane at room temperature for 5 min the RNA was cross-linked with an UV crosslinker as described above.

For *in vitro* transcription and hybridization to slot-blot membranes, chloroplasts from *Oenothera* leaves harvested 8-10 weeks after germination were isolated and counted according to a previously published protocol, applying the same minor modifications as described in Sobanski et al. (2019)^39^. Then, a chloroplast suspension containing 4.9 x 10^7^ chloroplasts was transferred to a fresh tube, centrifuged at 5,000 g for 1 min and the supernatant was removed. To start the *in vitro* transcription, 20 units of RNase Inhibitor (Promega GmbH), 50 μCi of [α-32P] UTP, and 94 μl Transcription Buffer (50 mM Tris—HCl pH 8.0, 10 mM MgCl_2_, 0.2 mM CTP, GTP, and ATP, 0.01 mM UTP, 10 mM 2-mercaptoethanol) were added, mixed and incubated for 10 min at 25°C. Next, the reaction was stopped by adding 10 μl of Stop Buffer [5% (w/v) Na-lauroylsarcosine, 50 mM Tris—HCl, pH 8.0, 25 mM EDTA] followed by a RNA isolation protocol, where 100 μl of phenol/chloroform/isoamylalcohol (25:24:1) was added to the reaction, vortexed, incubated for 10 min at room temperature, and centrifuged at 18,000 g for 10 min at 4°C. Afterwards, the upper phase was collected and nucleic acids were predicated overnight at −20°C using 3 volumes of 100% (v/v) ethanol, 0.3 M sodium acetate and 1 μl GlycoBlue™ (Invitrogen). On the next day, the sample was centrifuged at 20,000 g for 1 h at 4°C. After centrifugation, the pellet was washed in 75% (v/v) ethanol and dissolved in 50 μl of RNase-free water. Next, the RNA was denatured at 75°C for 15 min and cooled for 2 min on ice. Before hybridizing the slot-blots with the isolated RNA, the membrane was pre-hybridized with 20 ml of Church Buffer [1 mM EDTA, 7% (w/v) SDS, 0.5 M NaHPO4 pH 7.2] in hybridization tubes at 65°C for 1 h. Hybridization was performed at 65°C overnight. Subsequently, the membrane was washed once with 1 x SSC and 0.2% (w/v) SDS for 10 min, and once with 0.5 x SSC and 0.2% (w/v) SDS for 10 min. After washing, the membrane was wrapped in a transparent foil and exposed to a storage phosphor screen for 5 days. The signals were detected using an Amersham Typhoon IP scanner.

### 5’-RACE to map transcription start sites

TAP transcript 5’-end mapping in *Oenothera* chloroplasts was performed as previously described^87^. In brief, primary transcripts of bacteria and cell organelles harbor triphosphates at their 5’ ends, while processed transcripts possess monophosphates at this position. The TAP enzyme (tobacco acid pyrophosphatase) removes the additional phosphates from the 5’-end of primary transcripts. After this treatment, both primary and processed transcripts can serve as substrate for RNA ligase. This allows to distinguish between primary and processed transcripts, when +TAP and -TAP treated samples are compared. In -TAP samples, ligation products originating form primary transcripts are absent. Hence, to map the transcription start sites of the *psbB* operon and to distinguish them from processing sites in close proximity, a 5’-RACE from +TAP (Epicenter) and -TAP RNA samples was performed. For this, RNA samples of both treatments were ligated to an RNA linker (5’-GUGAUCCAACCGACGCGACAAGCUAAUGCAAGANNN-3’). After cDNA synthesis with a *psbB* gene-specific primer (psbB_cDNA_jn 5’-GCTGGCTGTCCATATAATGCATACAGC-3’), two PCRs were performed: The first PCR employed the linker-specific primer RUMSH1 (5’-TGATCCAACCGACGCGAC-3’) and the *psbB*-specific primer psbB_cDNA_jn. The second PCR used the linker-specific nested primer RUMSH2 (5’ ACCGACGCGACAAGCTAATGC-3’) and the primer psbB_5prime_jn (5’-GGAAAGGGATTTTAGGCATACCAATCG-3’). PCR products (30 μl of PCR solution) were run on 1% agarose gels (w/v) and, prior to sequencing, cloned into pCR^®^2.1-TOPO^®^ (Invitrogen).

### Statistical analysis

All numerical results are reported as mean ±SD. Statistical significance of the difference between experimental groups was analyzed by unpaired t-test using GraphPad Prism software. Differences were considered statistically significant for *P* < 0.05 or P < 0.01. Northern/western blots and run-on analyses were repeated at least twice. Representative data are shown.

## Data availability

Data supporting the findings of this manuscript are available from the corresponding authors upon reasonable request. A reporting summary for this Article is available as a Supplementary Information file. Sequence information has been deposited in GenBank with accession codes listed in the relevant tables and text passages. Source data underlying the association mapping are provided as Datasets S1 to S3.

## Acknowledgment

We thank Werner Dietrich and Wilfried Stubbe, who compiled the comprehensive *Oenothera* collection that allowed us to use the original material of the Renner school and Ralph E. Cleland. The Ornamental Plant Germplasm Centre is acknowledged for providing the line chicaginensis de Vries. We wish to thank Mario Anger for artwork, Christina Hedderich for help with the plastome sequences, Jennifer Jammrath for support with transcription start site mapping, Jörg Meurer for providing the Pbf1 antibody, Liliya Yaneva-Roder, and Wolfram Thiele for technical assistance, and the Max Planck Institute of Molecular Plant Physiology GreenTeam for nursing our material. This research was supported by the Max Planck Society (S.G., M.A.S., and R.B.).

## Authors contributions

A.Z., D.K., M.A.S., J.N., F.G., I.M. and S.G. performed the experimental work. All authors analyzed and discussed data. S.G. designed the study and wrote the manuscript. R.B. participated in writing.

### Materials & Correspondence

Correspondence and requests for materials should be addressed to S.G.

## Supplementary Information for

### Other supplementary materials for this manuscript include the following

Datasets S1 to S3

- S1: alignment_46_Oenothera_plastomes.fas
- S2: alignment_46_Oenothera_plastomes.sqd
- S3: consensus_46_Oenothera_plastomes_annotation.gb

### Reporting Summary fileSupplementary Text

#### Photosynthetic phenotype of AB-I plants

To understand the yellow-green *(lutescent)* AB-I phenotype (Fig. 2a), we performed a detailed characterization of its photosynthetic parameters. It appeared that the photosynthetic apparatus of *Oenothera* AB-I plants suffers from light-dependent damage. When plants were grown under three different light intensities (300, 450 and 600 μE m^-2^ s^-1^), no damage to the photosynthetic apparatus of AB-I occurred at 300 μE m^-2^ s^-1^ (see below). At 600 μE m^-2^ s^-1^, however, destruction of the photosynthetic apparatus was already so massive that it precluded a detailed photosynthetic characterization. We, therefore, characterized compatible AB-II and incompatible AB-I plants grown at 300 and 450 μE m^-2^ s^-1^, designated as low light (LL) and high light (HL), respectively, where pronounced differences were observed. Under LL, function and composition of the photosynthetic apparatus were indistinguishable between AB-II and AB-I. As judged from photosynthetic complex quantification by difference absorbance measurements normalized to a leaf area basis, the contents of photosystem II (PSII), cytochrome *b*_6_*f* complex (Cyt*b*_6_*f*), the mobile redox carrier plastocyanin (PC) and photosystem I (PSI) were indistinguishable between the genotypes (Fig. 2b). Also, the maximum quantum efficiency of PSII in the dark-adapted state (F_V_/F_M_) was identical (0.79), clearly showing that no photoinhibition of PSII occurred under these conditions. The same was true for the total chlorophyll content per leaf area (Supplementary Fig. 1a). By contrast, the increase in light intensity to 450 μE m^-2^ s^-1^ resulted in drastic changes in photosynthetic complex accumulation, F_V_/F_M_ and chlorophyll content in the two genotypes. AB-II plants responded to the increased growth light intensity by strongly increasing their chlorophyll content and the contents of all redox-active components of the electron transport chain, ranging from a more than two-fold increase of PSII content to a 50% increase in PSI content (Fig. 2b). These increases represent the typical light acclimation response that occurs when plants previously grown under light-limited conditions are transferred to a higher light intensities^1^. AB-I plants were incapable of performing this light acclimation response efficiently. Their PSII content increased only by 40%, and the strong decrease in F_V_/F_M_ suggested that a substantial number of PSII centers were photodamaged. Also, contents of Cytb_6_*f* complex and PC increased to a much lesser degree than in AB-II, and PSI content remained essentially unaltered (Fig. 2b). In line with these observations, the chlorophyll content per leaf area increased only by 25% in AB-I (Supplementary Fig. 1a).

To assess possible consequences of these different light acclimation responses of AB-II and AB-I on the relative antenna cross sections of the two photosystems, chlorophyll-a fluorescence emission spectra at 77K were recorded (Supplementary Fig. 1b). For better comparability, the spectra were normalized to the PSII emission maximum at 687 nm wavelength. At 300 μE m^-2^ s^-1^, AB-II had a higher PSI emission signal than AB-I, in line with its slightly higher ratio of PSI to PSII. With increasing light intensity, the photosystem I/light harvesting complex I (PSI-LHCI) emission signal, peaking at 733 nm wavelength, decreased in both AB-II and AB-I, well in line with the more pronounced increase in PSII content in both genotypes. No indications for the presence of free, uncoupled light-harvesting complex I (LHCI) or light-harvesting complex II (LHCII) were observed. These would be expected to result in additional emission signals at 680 nm wavelength (indicative of free LHCII), or between 705 and 730 nm wavelength (indicative of the presence of uncoupled LHCI)^2–4^. Therefore, the decreased F_V_/F_M_ ratio of AB-I at 450 μE m^-2^ s^-1^ (Fig. 2b) cannot be explained by the presence of uncoupled antenna, but has to be attributed to photoinhibition of the PSII reaction centers themselves.

Finally, chloroplast ATP synthase (ATPase) activity and accumulation were assessed (Supplementary Fig. 1c,d). Dark-interval relaxation kinetics of the electrochromic shift, a measure for the proton motive force across the thylakoid membrane, were used to determine the thylakoid conductivity for protons (gH^+^). The latter, in turn, is a proxy for ATPase activity^5,6^. Interestingly, as judged from western blot analyses of the AtpA protein, a core subunit of the ATP synthase complex, a slight reduction of ATP synthase content might be present in AB-I plants, at least under low light conditions (Supplementary Fig. 1d). This, however, did not lead to significant differences in ATP synthase activity between the genotypes (Supplementary Fig. 1c). Moreover, we did not observe significant differences between the growth light intensities. In this context, it should be mentioned that, in contrast to the complex quantifications expressed on a leaf area basis, gH^+^ is a measure for ATP synthase activity per thylakoid membrane and not per leaf area. Hence, since chlorophyll content increased with light intensity, it is likely that also total ATP synthase activity per leaf area was higher in plants grown at 450 μE m^-2^ s^-1^ compared to plants grown at 300 μE m^-2^ s^-1^. Nonetheless, this effect is independent of the compatible/incompatible situation of AB-II/AB-I plants.

In summary, AB-I plants display a light-dependent photosynthesis phenotype that cannot be attributed to a single component of the electron transport chain. However, the phenotype is independent of ATP synthase and PC function.

### Northern blot analyses of *psbB* operon transcripts

Transcript analysis of the *psbB* operon showed a clear decrease in mRNA accumulation in AB-I plants under HL conditions, whereas no differences were detected for AB-I plants in LL (Supplementary Fig. 2a,c). Upon hybridization with a *psbB* probe, four major psbB-containing transcript species were detected. The pentacistronic *psbB-psbT-psbH-petB-petD* transcript (5.6 kb with the introns of *petB* and *petD,* and 3.9 kb without these introns), the tricistronic *psbB-psbT-psbH* transcript (2.6 kb) and the dicistronic mature *psbB-psbT* transcript (1.8 kb). For details on the maturation of the *psbB* operon, see Westhoff and Herrman (1988)^7^. Surprisingly, a processing defect was detected in AB-I plants under HL conditions in that the AB-I plants lack the 3.9 kb transcript with the *petB* and *petD* introns spliced out (Supplementary Fig. 2c, boxed in red). The same transcript species were detected when blots were hybridized to a *psbT* probe. Hybridization to a *psbH* probe revealed a down-regulation of all *psbH* transcripts. Similarly to hybridization with the *psbB* probe, virtual absence of the 3.9 kb transcripts was observed. For both *petB* and *petD,* a decrease of polycistronic and monocistronic transcripts was observed. Both *petB* and *petD* mRNAs appear to accumulate to higher levels in AB-I at LL. Again, absence of the 3.9 kb transcript speceis was confirmed.

In summary, the data show that expression of the entire *psbB* operon is affected by the deletion. In the absence of any polymorphism between plastome I and II in the whole operon (Supplementary Dataset 1-3), the observed processing defect is likely to be a secondary consequence of the deletion. The precise molecular mechanism underlying this effect is currently unknown.

### Supplementary Figures and Tables

**Supplementary Fig. 1.**
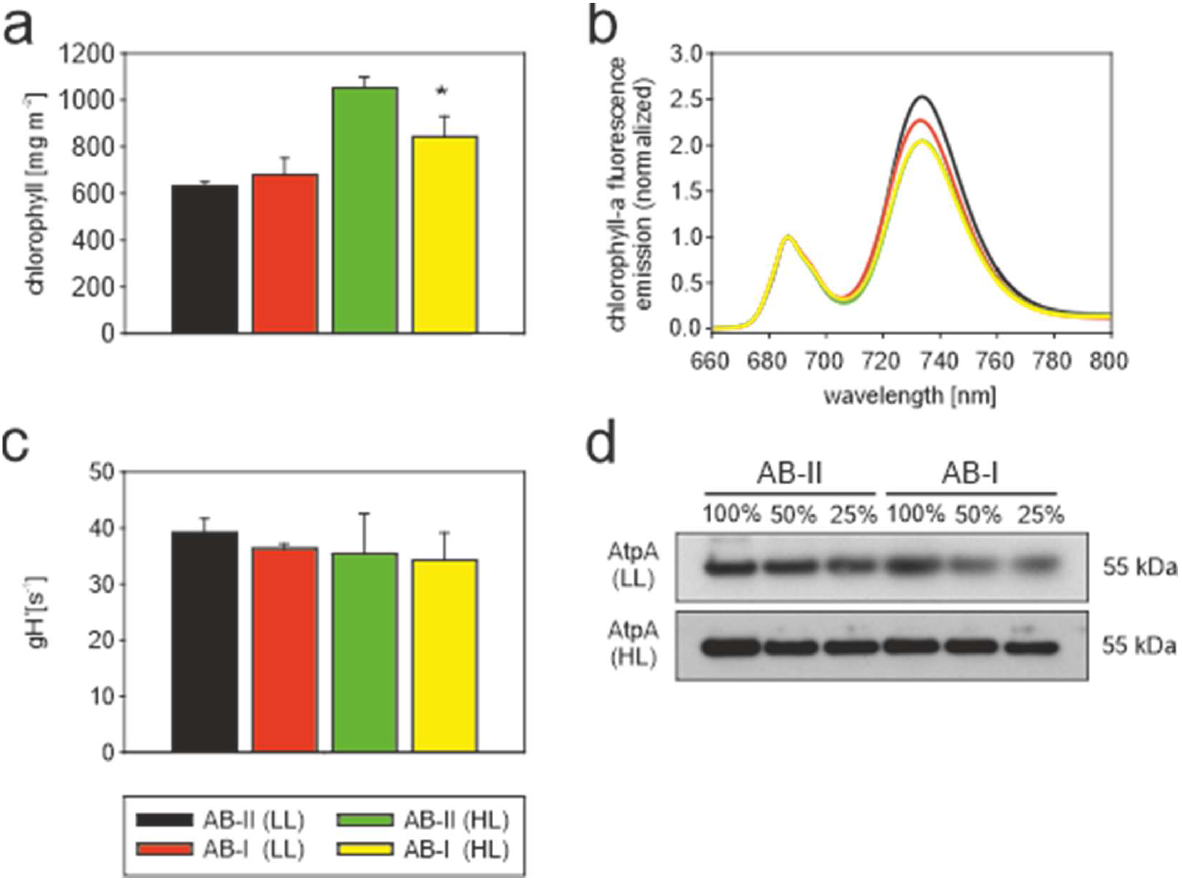
Photosynthetic parameters of compatible AB-II and incompatible AB-I plants grown under LL or HL conditions. **a**, Chlorophyll content per leaf area. Bars represent mean values ±SD (n = 4-6). Asterisk indicates significant difference from AB-II HL t-test, P<0.05 (t = 5.72, df = 8). **b**, Chlorophyll-a fluorescence emission spectra at 77K. Bars represent mean values ±SD (n = 4-6).**c**, Thylakoid membrane conductivity for protons (gH^+^) as proxy for ATP synthase activity. **d**, Western blot analysis of the AtpA protein, a core subunit of the ATP synthase complex. Samples are normalized to chlorophyll content. This experiment was performed three times independently with similar results.

**Supplementary Fig. 2.**
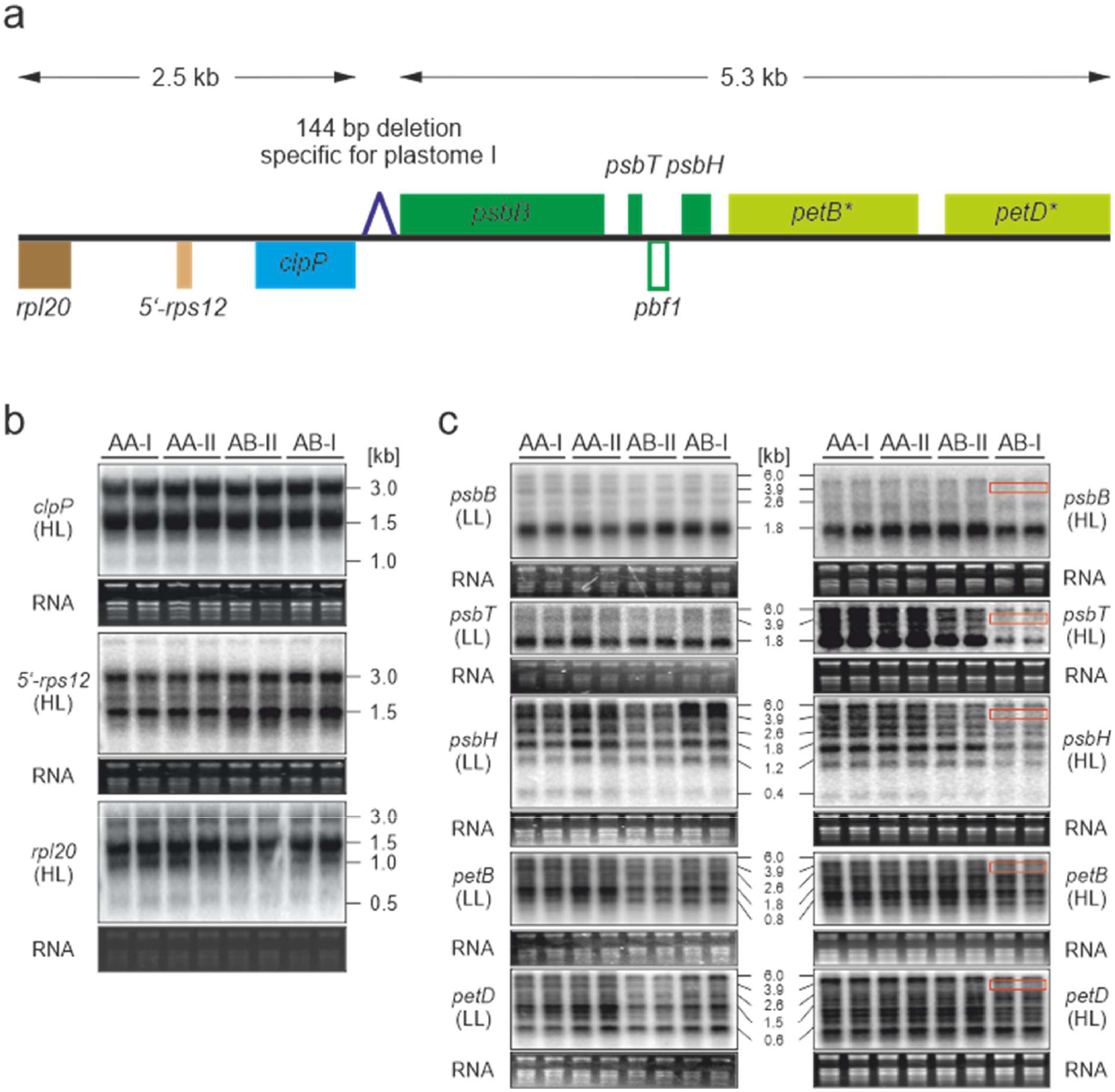
Northern blot analyses of the *clpP* and *psbB* operons in compatible (AA-I, AA-II, AB-II) and incompatible material (AB-I). **a**, Sequence context and position of the deletion specific to plastome I. Note that *clpP* is co-transcribed with exon 1 of the trans-spliced gene *rps12 (5’-rps12)* and with *rpl20.* The intron-containing genes *petB* and *petD* are marked by asterisks. **b**, Northern blot analyses of *clpP* operon transcripts in plants grown under HL conditions. Northern blots were performed two times independently with similar results. **c**, Northern blot analyses of *psbB* operon transcripts under LL and HL conditions. The processing defect is boxed in red. These experiments were performed two times independently with similar results.

**Supplementary Table 1.**
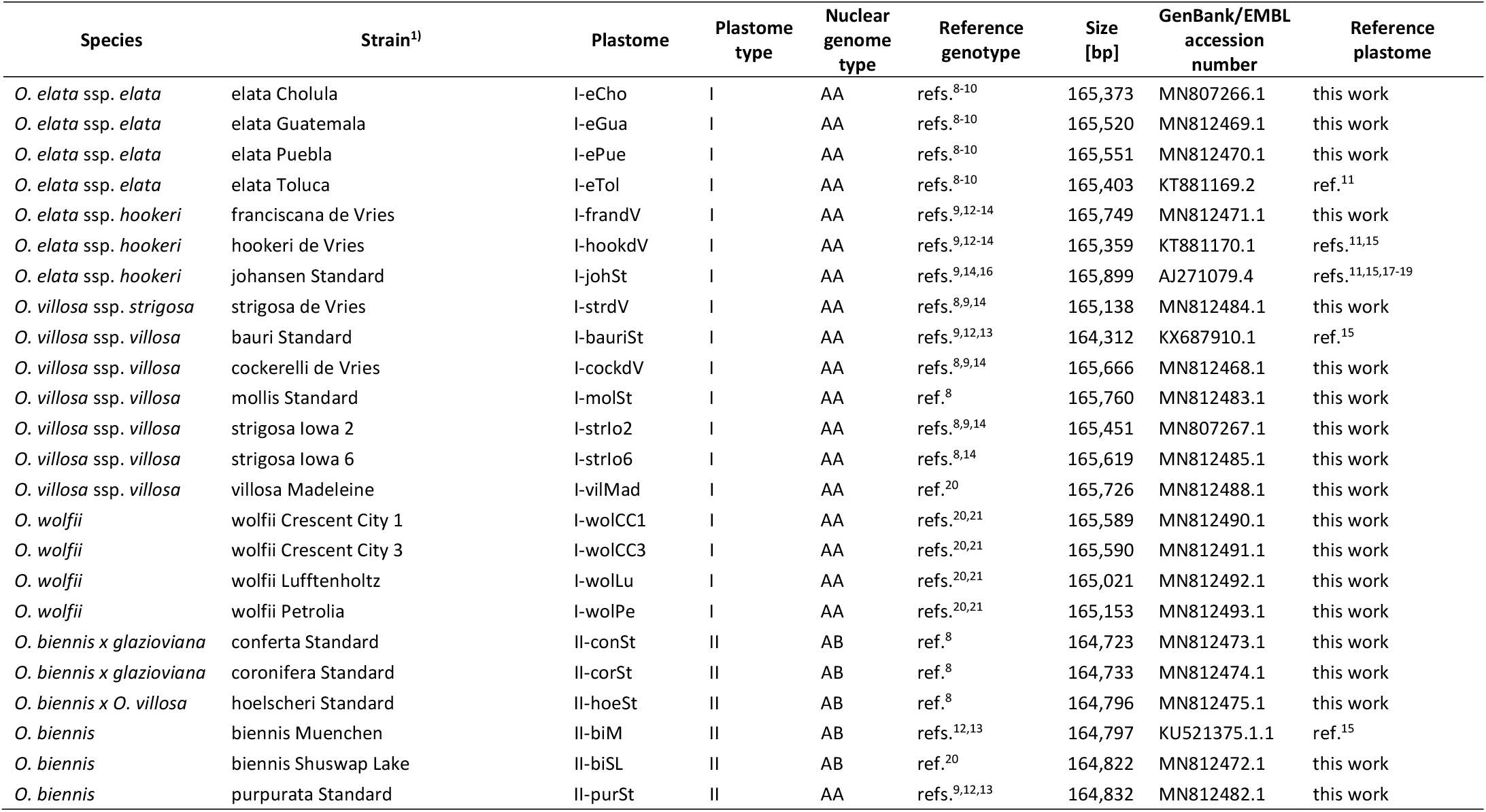
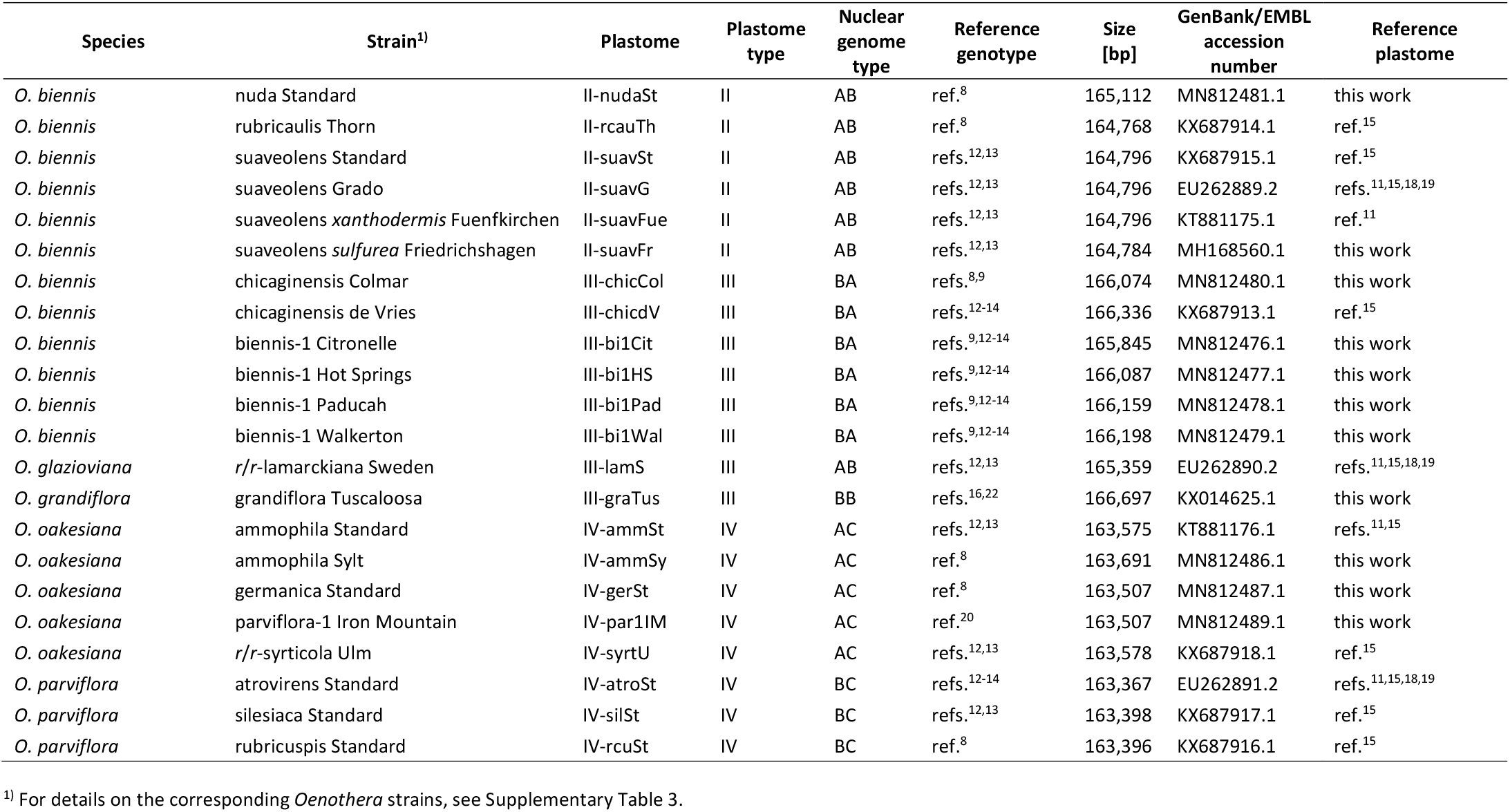
Accession number, genome size, genetic information, and corresponding nuclear genotype of *Oenothera* plastomes employed for association mapping.

**Supplementary Table 2.**
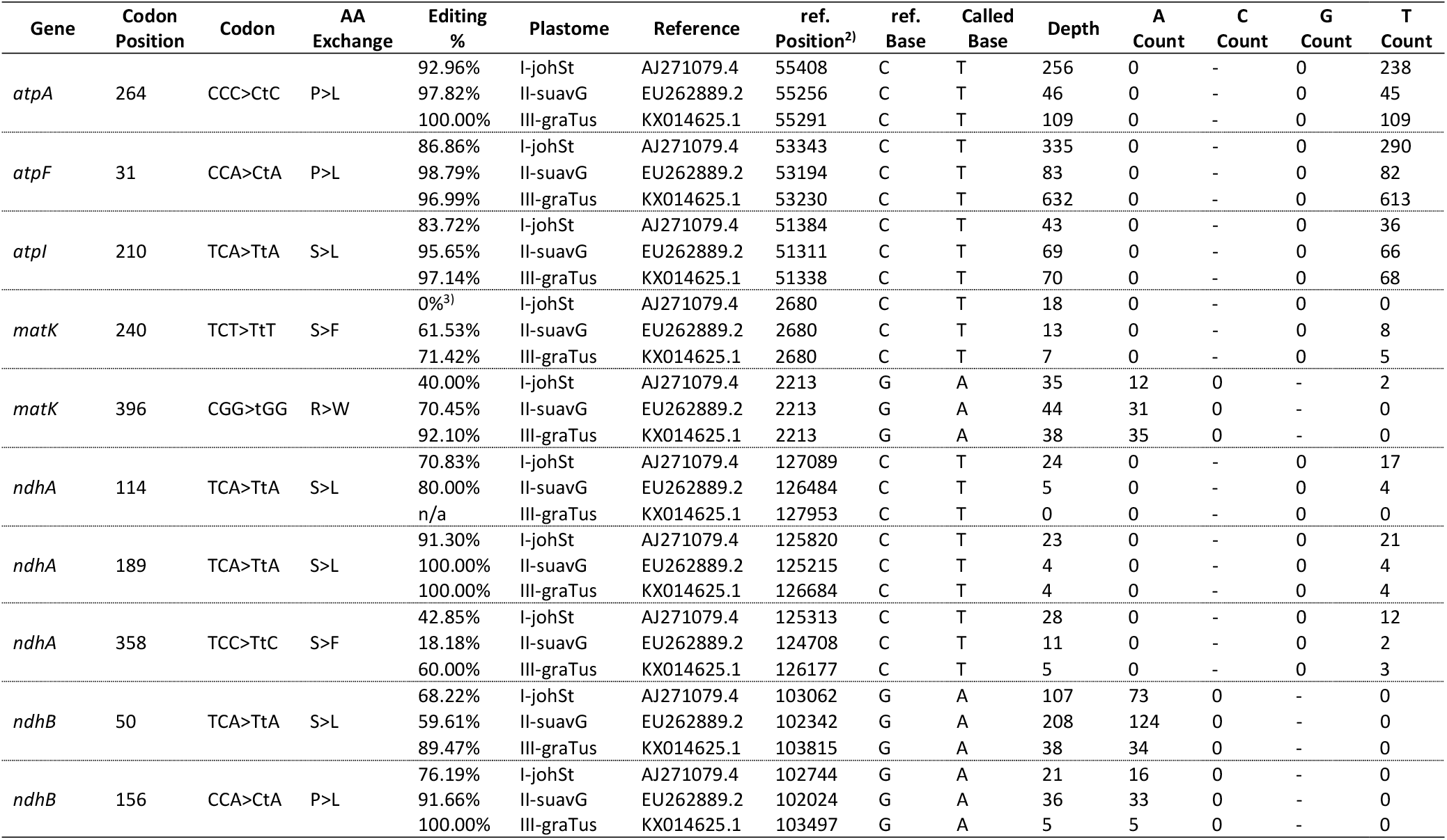
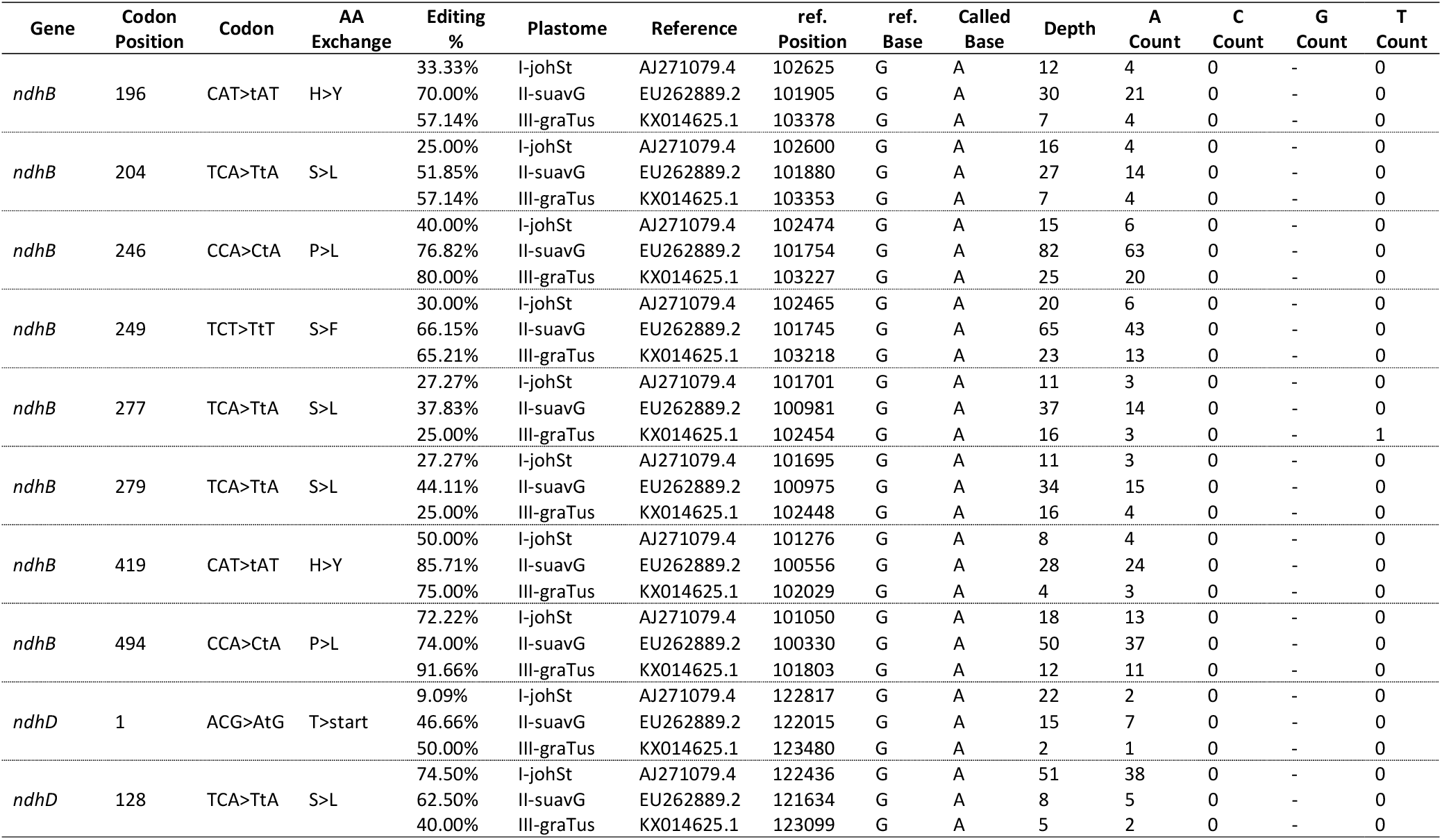
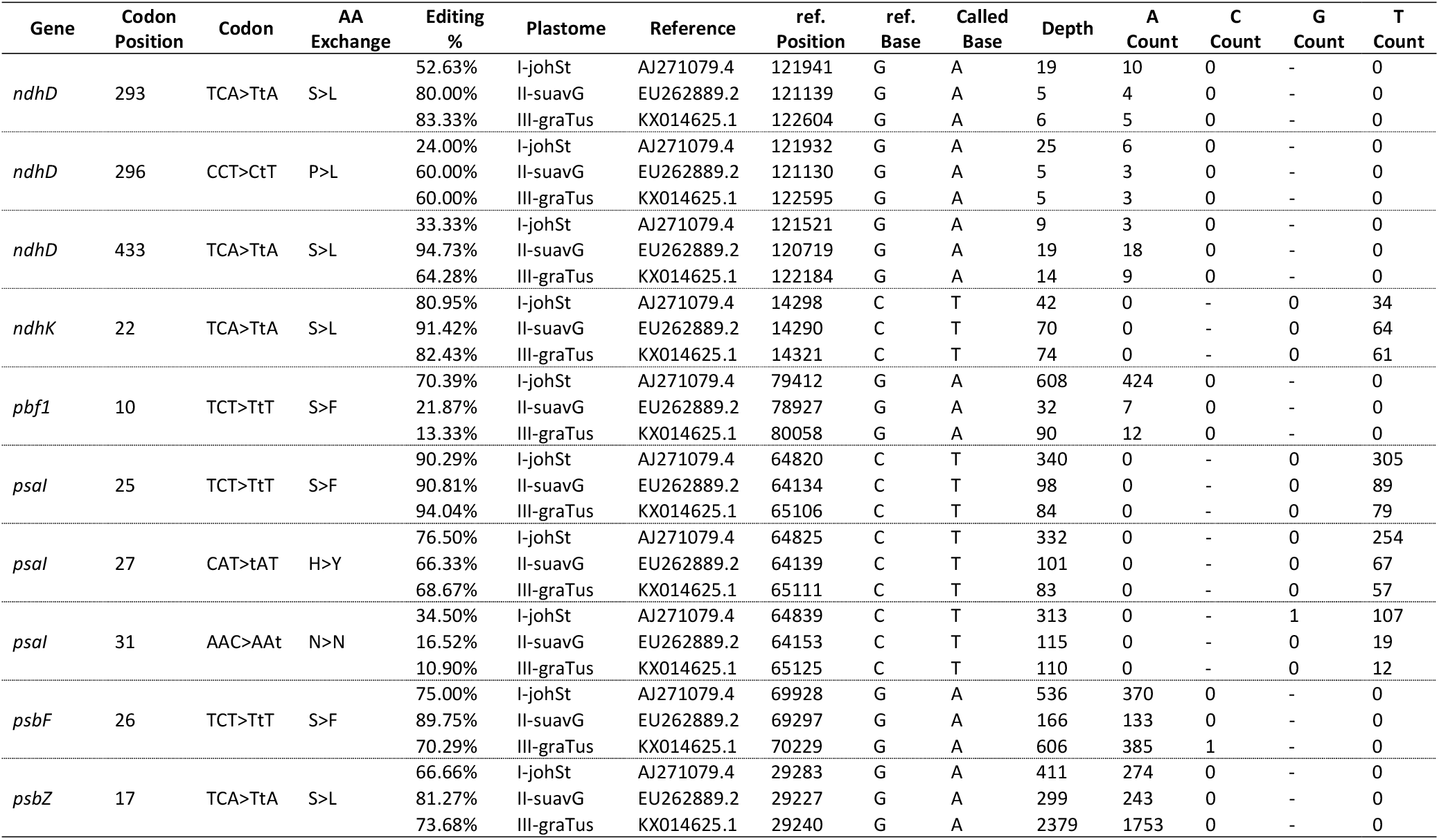
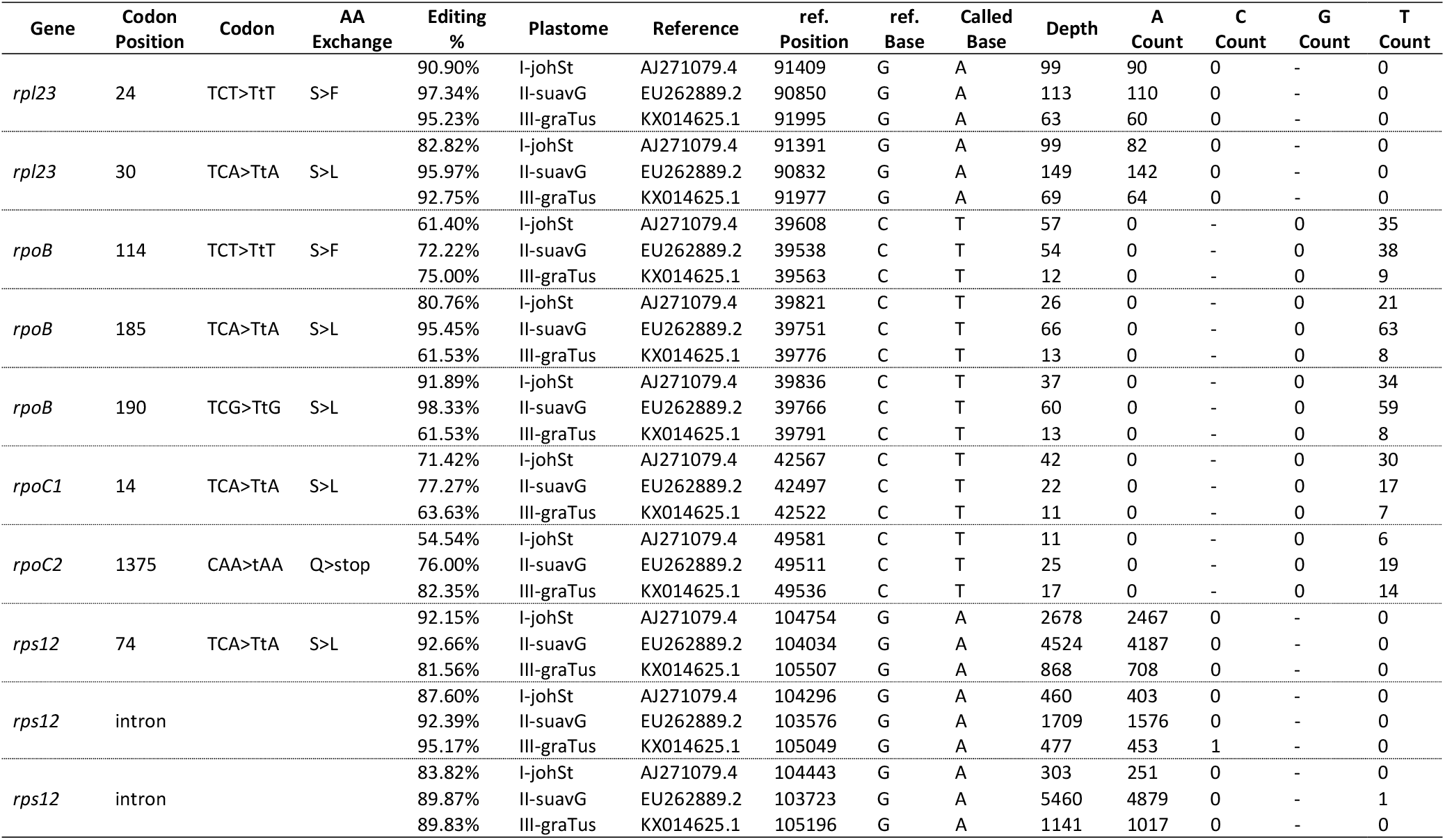
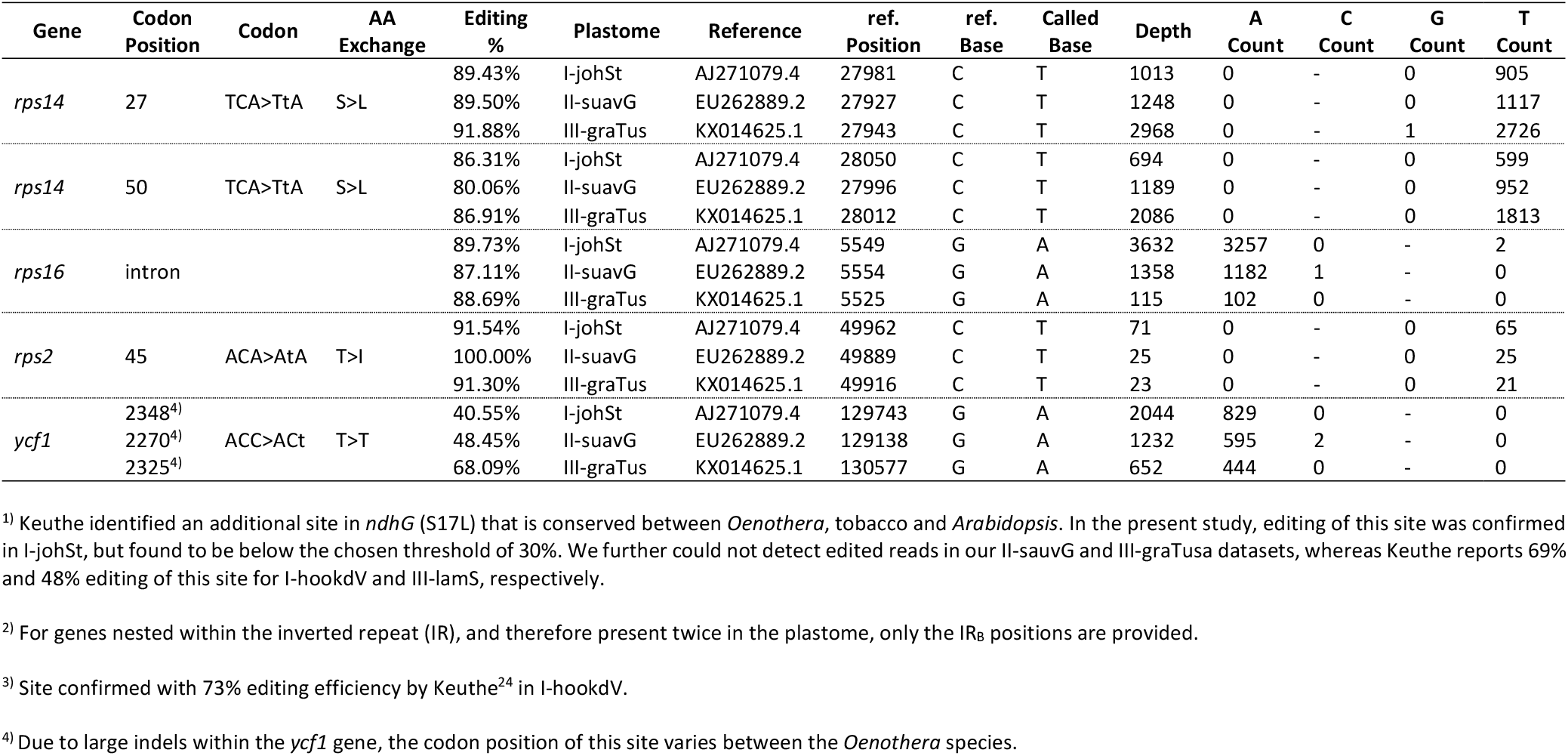
Chloroplast mRNA editotype and cDNA mapping results of three *Oenothera* species from subsection *Oenothera.* For details, see Methods. See refs.^23,24^ for additional data^1)^.

**Supplementary Table 3.** Origin and collector’s information of the *Oenothera* strains used in this work.

| Species | Strain | Locality | Collection date | Collector | Reference |
| --- | --- | --- | --- | --- | --- |
| *O. elata* ssp. *elata* | elata Cholula | Mexico, Puebla, 6 miles north-west of Cholula de Rivadavia | before 1949 | P. A. Munz | refs.<sup>25,26</sup> |
| *O. elata* ssp. *elata* | elata Guatemala | Guatemala, Guatemala, Guatemala City | 1945 | P. Weatherwax | ref.<sup>27</sup> |
| *O. elata* ssp. *elata* | elata Puebla | Mexico, Puebla, garden at Puebla | before 1949 | P. A. Munz | refs.<sup>25,26</sup> |
| *O. elata* ssp. *elata* | elata Toluca | Mexico, Mexico, garden at Toluca de Lerdo | 1937 | P. A. Munz | refs.<sup>25,27</sup> |
| *O. elata* ssp. *hookeri* | franciscana de Vries<sup>1)</sup> | USA, CA, Monterey Co., Carmel Beach | 1905 | C. P. Smith | refs.<sup>28,29</sup> |
| *O. elata* ssp. *hookeri* | hookeri de Vries | USA, CA, Alameda Co., near Berkeley | 1904 | H. de Vries | ref.<sup>30</sup> |
| *O. elata* ssp. *hookeri* | johansen Standard | USA, CA, Sutter Co., roadside between Nicholas and Yuba City | 1927 | C. B. Wolf | ref.<sup>31</sup> |
| *O. villosa* ssp. *strigosa* | strigosa de Vries | USA, WY, Park Co., Yellowstone National Park near Mammoth Hot Springs | 1904 | H. de Vries | ref.<sup>30</sup> |
| *O. villosa* ssp. *villosa* | bauri Standard | Poland, Kujawsko-Pomorskie, near Toruń | before 1942 | R. Hölscher | ref.<sup>32</sup> |
| *O. villosa* ssp. *villosa* | cockerelli de Vries | USA, CO, Boulder Co., near Boulder | 1905 | T. D. A. Cockerell | ref.<sup>30</sup> |
| *O. villosa* ssp. *villosa* | mollis Standard | Germany, Brandenburg, in sandy soil near Jüterbog | 1934 | O. Renner | ref.<sup>33</sup> |
| *O. villosa* ssp. *villosa* | strigosa Iowa 2 | USA, IA, Dickson Co., in a gravel pit one mile south-west of Manhattan Beach | 1930 | J. B. Eisen | ref.<sup>34</sup> |
| *O. villosa* ssp. *villosa* | strigosa Iowa 6 | USA, IA, Dickson Co., on the slope of a gravel pit one mile south-west of Manhattan Beach | 1930 | J. B. Eisen | ref.<sup>35</sup> |
| *O. villosa* ssp. *villosa* | villosa Madeleine | Canada, QC, Gaspésie-Îles-de-la-Madeleine Magdalen Islands | before 1967 | Anonymous | ref.<sup>20</sup> |
| *O. wolfii* | wolfii Crescent City 1 | USA, CA, Del Norte Co., Crescent City near marina | 1977 | P. C. Hoch | refs.<sup>20,21</sup> |
| *O. wolfii* | wolfii Crescent City 3 | USA, CA, Del Norte Co., Crescent City near marina | 1977 | P. C. Hoch | refs.<sup>20,21</sup> |
| *O. wolfii* | wolfii Lufftenholtz | USA, CA, Humboldt Co., Lufftenholtz, Beach County Park south of Trinidad | 1975 | J. D. Ackerman and A. M. Montalvo | refs.<sup>20,21</sup> |
| *O. wolfii* | wolfii Petrolia | USA, CA, Humboldt Co., Petrolia | 1977 | P. C. Hoch | ref.<sup>20</sup> |

Supplementary Table 3. (continued)
| Species | Strain | Locality | Collection date | Collector | Reference |
| --- | --- | --- | --- | --- | --- |
| *O. biennis x glazioviana* | conferta Standard | France, Calvados, near Cabourg north-east of Caen, directly on a costal sand dune | before/in 1942 | F. Hilpert | Rrefs.<sup>36</sup> |
| *O. biennis x glazioviana* | coronifera Standard | Germany, Brandenburg, railway embankment at Zinna Abbey near Jüterbog | 1936 | O. Renner | ref.<sup>37</sup> |
| *O. biennis x villosa* | hoelscheri Standard | Poland, Kuyavian-Pomeranian, near Vistula River at Włocławek | before 1942 | R. Hölscher | ref.<sup>38</sup> |
| *O. biennis* | biennis Muenchen | Germany, Bavaria, Munich, Nymphenburg Garden | 1914 | O. Renner | ref.<sup>39</sup> |
| *O. biennis* | biennis Shuswap Lake | Canada, BC, Shuswap Lake | 1977 | G. B. Straley | ref.<sup>20</sup> |
| *O. biennis* | nuda Standard | France, Isère, narrow-gauge railway embankment at both sides of Saint-Laurent-du-Pont | 1947 | A. Gagnieu | ref.<sup>38</sup> |
| *O. biennis* | purpurata Standard<sup>2)</sup> | chromosome translocation mutant of material reassembling *biennis cruciata* Klebahn<sup>3)</sup> | isolated in 1914 | H. Klebahn | refs.<sup>40,41</sup> |
| *O. biennis* | rubricaulis Thorn | Poland, Kujawsko-Pomorskie, Vistula River near Toruń | before 1941 | R. Hölscher | ref.<sup>42</sup> |
| *O. biennis* | suaveolens Standard | France, Seine-et-Marne, forest of Fontainebleau | 1912 | L. Blaringhem | ref.<sup>43</sup> |
| *O. biennis* | suaveolens Grado | Italy, Friuli-Venezia Giulia, dune near Grado at the Adriatic sea | before 1950 | H. Zeidler | refs.<sup>44,45</sup> |
| *O. biennis* | suaveolens *xanthodermis* Fuenfkirchen | Hungary, Baranya, near Pécs | before 1949 | E. Preuss | refs.<sup>44,45</sup> |
| *O. biennis* | suaveolens *sulfurea* Friedrichshagen | Germany, Berlin, Treptow-Köpenick, at railway station Friedrichshagen-Hirschgarten | 1937 | O. Renner | refs.<sup>44,45</sup> |
| *O. biennis* | chicaginensis Colmar | France, Haut-Rhin, fallow on the road between Niederhergheim and mill at Dessenheim | 1943 | E. Issler | ref.<sup>38</sup> |
| *O. biennis* | chicaginensis de Vries | USA, IL, Cook Co., Chicago, near Jackson Park | 1904 | H. de Vries | ref.<sup>30</sup> |
| *O. biennis* | biennis-1 Citronelle | USA, AL, Mobile Co., Citronelle | 1935 | P. A. Munz | ref.<sup>46</sup> |
| *O. biennis* | biennis-1 Hot Springs | USA, AR, Garland Co., Hot Springs | before 1958 | Anonymous | ref.<sup>47</sup> |

**Supplementary Table 3.** (continued)
| Species | Strain | Locality | Collection date | Collector | Reference |
| --- | --- | --- | --- | --- | --- |
| *O. biennis* | biennis-1 Paducah | USA, KY, McCracken Co., eleven miles west of Paducah | 1935 | P. A. Munz | ref.<sup>34</sup> |
| *O. biennis* | biennis-1 Walkerton | USA, IN, St. Joseph Co., Walkerton | before 1958 | Anonymous | ref.<sup>47</sup> |
| *O. glazioviana* | *r/r-lamarckiana* Sweden | Sweden, Skåne Län, garden in Almaröd | 1906 | N. Heribert-Nilsson | ref.<sup>48</sup> |
| *O. grandiflora* | grandiflora Tuscaloosa | USA, AL, Mobile Delta | 1944 | J. S. Lloyd | ref.<sup>49</sup> |
| *O. oakesiana* | ammophila Standard | Germany, Schleswig-Holstein, Helgoland | 1922 | E. Hoepfener | ref.<sup>50</sup> |
| *O. oakesiana* | ammophila Sylt | Germany, Schleswig-Holstein, southern tip of the island of Sylt | before 1963 | Anonymous | ref.<sup>8</sup> |
| *O. oakesiana* | germanica Standard | Germany, Berlin, Berlin-Rahnsdorf | 1918 | E. Baur | refs.<sup>50,51</sup> |
| *O. oakesiana* | parviflora-1 Iron Mountain<sup>4)</sup> | USA, MI, Dickinson Co., five miles south-east of Iron Mountain | 1938 | P. A. Munz | ref.<sup>52</sup> |
| *O. oakesiana* | *r/r-syrticola* Ulm | Germany, Baden-Württemberg, Danube River near Ulm | 1917 | O. Renner | ref.<sup>33</sup> |
| *O. parviflora* | atrovirens Standard<sup>5)</sup> | USA, NY, Erie Co., Sandy Hill near Lake George | 1902/1903 | D. T. MacDouglas | refs.<sup>30,53</sup> |
| *O. parviflora* | silesiaca Standard | Poland, Dolnośląskie, bank of Bóbr River near Nowogrodziec | 1937 | O. Renner | ref.<sup>42</sup> |
| *O. parviflora* | rubricuspis Standard | Germany, Hessen, railway embankment between Neu-Isenburg and Luisa near Frankfurt on the Main | 1942/1943 | O. Burck, F. Laibach and E. Fischer | ref.<sup>45</sup> |

<sup>1)</sup> The strain franciscana de Vries is a derivative of Davis's franciscana B<sup>54</sup>, as summarized in Davis (1916)<sup>55</sup>.
<sup>2)</sup> Originally described by Klebahn as *Oenothera biennis rubicalyx*<sup>40</sup>.
<sup>3)</sup> Derivative of parent plant No. 347, similar to biennis *cruciata* Klebahn collected in Population 4 near Bad Bevensen (Germany, Niedersachsen)<sup>40</sup>.
<sup>4)</sup> This line was originally described as parviflora-1 (= BC-IV = *O. parviflora*) by Cleland<sup>56</sup>, but identified as AC-IV (= parviflora-2 = *O. oakesiana*) by Wasmund<sup>20</sup>.
<sup>5)</sup> According to Renner<sup>53</sup> this line was originally "received from Amsterdam" by N. v. Gescher in 1907. The material is quite likely identical to that collected by D. T. MacDouglas in 1902/1903 as described in de Vries (1913)<sup>30</sup>. Also see Bartlett (1914)<sup>57</sup>.

**Supplementary Table 4.** Chloroplast substitution lines and corresponding wild types used in this work.

| Line and genotype | Plastome | Nuclear genome<sup>1)</sup> | Chloroplast donor strain<sup>2)</sup> | Nucleus donor strain<sup>2)</sup> | Phenotype | Produced by | Reference |
| --- | --- | --- | --- | --- | --- | --- | --- |
| AA-I | I-johSt | *<sup>h</sup>johansen Standard·<sup>h</sup>johansen Standard* | johansen Standard | johansen Standard | green | wild type | refs.<sup>9,14,16,31</sup> |
| AA-II | II-suavG | *<sup>h</sup>johansen Standard·<sup>h</sup>johansen Standard* | suaveolens Grado | johansen Standard | green | W. Stubbe | refs.<sup>11,58</sup>, this work |
| AB-I | I-johSt | *<sup>G</sup>albicans·<sup>G</sup>flavens* | johansen Standard | suaveolens Grado | *lutescent* | S. Greiner | this work |
| AB-II | II-suavG | *<sup>G</sup>albicans·<sup>G</sup>flavens* | suaveolens Grado | suaveolens Grado | green | wild type | refs.<sup>12,13,44</sup> |

<sup>1)</sup> See Material and Methods for details.
<sup>2)</sup> For details on the donor stains, see Supplementary Table 3

**Supplementary Table 5.**
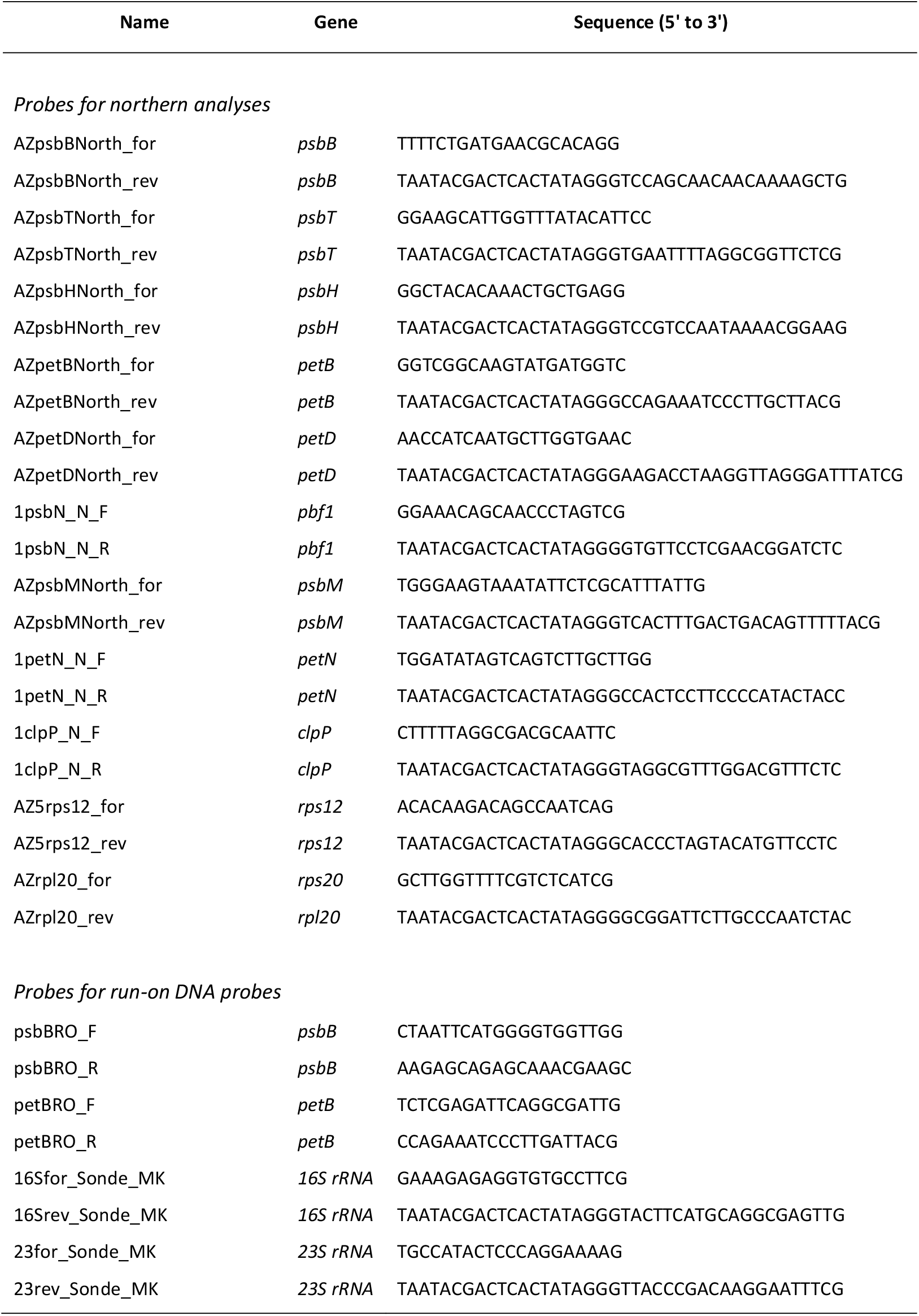
Oligonucleotides used for the generation of probes for northern blot and run-on transcription analyses.

## References

1 Burton, R. S., Pereira, R. J. & Barreto, F. S. Cytonuclear genomic interactions and hybrid breakdown. Annual Review of Ecology, Evolution, and Systematics 44, 281–302, doi:10.1146/annurev-ecolsys-110512-135758 (2013).

2 Chou, J.-Y. & Leu, J.-Y. Speciation through cytonuclear incompatibility: insights from yeast and implications for higher eukaryotes. Bioessays 32, 401–411 (2010).

3 Fishman, L. & Sweigart, A. L. When two rights make a wrong: the evolutionary genetics of plant hybrid incompatibilities. Annu. Rev. Plant Biol. 69, 707–731, doi:10.1146/annurev-arplant-042817-040113 (2018).

4 Greiner, S. & Bock, R. Tuning a ménage à trois: co-evolution and co-adaptation of nuclear and organellar genomes in plants. Bioessays 35, 354–365 (2013).

5 Levin, D. A. The cytoplasmic factor in plant speciation. Systematic Botany 28, 5–11 (2003).

6 Bogdanova, V. S. Genetic and molecular genetic basis of nuclear-plastid incompatibilities. Plants 9, 23 (2020).

7 Postel, Z. & Touzet, P. Cytonuclear genetic incompatibilities in plant speciation. Plants 9, 487 (2020).

8 Greiner, S., Rauwolf, U., Meurer, J. & Herrmann, R. G. The role of plastids in plant speciation. Molecular Ecology 20, 671–691 (2011).

9 Schmitz-Linneweber, C. et al. Pigment deficiency in nightshade/tobacco cybrids is caused by the failure to edit the plastid ATPase alpha-subunit mRNA. Plant Cell 17, 1815–1828 (2005).

10 Harrison, J. S. & Burton, R. S. Tracing hybrid incompatibilities to single amino acid substitutions. Mol Biol Evol 23, 559–564 (2006).

11 Lee, H.-Y. et al. Incompatibility of nuclear and mitochondrial genomes causes hybrid sterility between two yeast species. Cell 135, 1065–1073 (2008).

12 Meiklejohn, C. D. et al. An incompatibility between a mitochondrial tRNA and its nuclear-encoded tRNA synthetase compromises development and fitness in *Drosophila*. PLOS Genetics 9, e1003238, doi:10.1371/journal.pgen.1003238 (2013).

13 Barnard-Kubow, K. B., So, N. & Galloway, L. F. Cytonuclear incompatibility contributes to the early stages of speciation. Evolution 70, 2752–2766, doi:10.1111/evo.13075 (2016).

14 Fishman, L. & Willis, J. H. A cytonuclear incompatibility causes anther sterility in *Mimulus* hybrids. Evolution 60, 1372–1381 (2006).

15 Stubbe, W. The role of the plastome in evolution of the genus *Oenothera*. Genetica 35, 28–33 (1964).

16 Dietrich, W., Wagner, W. L. & Raven, P. H. Systematics of Oenothera section Oenothera subsection Oenothera (Onagraceae). 1st edn, (The American Society of Plant Taxonomists, 1997).

17 Greiner, S. & Köhl, K. Growing evening primroses (Oenothera). Front. Plant Sci. 5, 38, doi:10.3389/fpls.2014.00038 (2014).

18 Stubbe, W. & Raven, P. H. A genetic contribution to the taxonomy of *Oenothera* sect. *Oenothera* (including subsection *Euoenothera, Emersonia, Raimannia* and *Munzia)*. Plant Systematics and Evolution 133, 39–59 (1979).

19 Hollister, J. D., Greiner, S., Johnson, M. T. J. & Wright, S. I. Hybridization and a loss of sex shape genome-wide diversity and the origin of species in the evening primroses (Oenothera, Onagraceae). New Phytologist 224, 1372–1380, doi:10.1111/nph.16053 (2019).

20 Cleland, R. E. Chromosome structure in *Oenothera* and its effect on the evolution of the genus. Cytologia 22, 5–19 (1957).

21 Cleland, R. E. Oenothera – Cytogenetics and Evolution. 1st edn, (Academic Press Inc., 1972).

22 Rauwolf, U., Golczyk, H., Meurer, J., Herrmann, R. G. & Greiner, S. Molecular marker systems for *Oenothera* genetics. Genetics 180, 1289–1306, doi:10.1534/genetics.108.091249 (2008).

23 Golczyk, H., Massouh, A. & Greiner, S. Translocations of chromosome end-segments and facultative heterochromatin promote meiotic ring formation in evening primroses. Plant Cell 26, 1280–1293, doi:10.1105/tpc.114.122655 (2014).

24 Raven, P. H., Dietrich, W. & Stubbe, W. An outline of the systematics of *Oenothera* subsect. *Euoenothera* (Onagraceae). Systematic Botany 4, 242–252 (1979).

25 Arntz, M. A. & Delph, L. F. Pattern and process: evidence for the evolution of photosynthetic traits in natural populations. Oecologia 127, 455–467 (2001).

26 Flood, P. J. Using natural variation to understand the evolutionary pressures on plant photosynthesis. Current Opinion in Plant Biology 49, 68–73, doi:https://doi.org/10.1016/j.pbi.2019.06.001 (2019).

27 Greiner, S. et al. The complete nucleotide sequences of the five genetically distinct plastid genomes of *Oenothera*, subsection *Oenothera:* I. Sequence evaluation and plastome evolution. Nucleic Acids Research 36, 2366–2378, doi:10.1093/nar/gkn081 (2008).

28 Hollister, J. D. et al. Recurrent loss of sex is associated with accumulation of deleterious mutations in *Oenothera*. Molecular Biology and Evolution 32, 896–905, doi:10.1093/molbev/msu345 (2015).

29 Levy, M. & Levin, D. A. Genic heterozygosity and variation in permanent translocation heterozygotes of the *Oenothera biennis* complex. Genetics 79, 493–512 (1975).

30 Anstett, D. N. et al. Can genetically based clines in plant defence explain greater herbivory at higher latitudes? Ecology Letters, n/a–n/a, doi:10.1111/ele.12532 (2015).

31 Greiner, S. et al. The complete nucleotide sequences of the 5 genetically distinct plastid genomes of *Oenothera*, subsection *Oenothera:* II. A microevolutionary view using bioinformatics and formal genetic data. Molecular Biology and Evolution 25, 2019–2030, doi:10.1093/molbev/msn149 (2008).

32 Schottler, M. A. & Tóth, S. Z. Photosynthetic complex stoichiometry dynamics in higher plants: environmental acclimation and photosynthetic flux control. Front. Plant Sci. 5, 188–188, doi:10.3389/fpls.2014.00188 (2014).

33 Cleland, R. E. Cyto-taxonomic studies on certain Oenotheras from California. Proc. Am. Philos. Soc. 75, 339–429 (1935).

34 Kahlau, S., Aspinall, S., Gray, J. & Bock, R. Sequence of the tomato chloroplast DNA and evolutionary comparison of Solanaceous plastid genomes. Journal of Molecular Evolution 63, 194–207, doi:10.1007/s00239-005-0254-5 (2006).

35 Takenaka, M., Zehrmann, A., Verbitskiy, D., Härtel, B. & Brennicke, A. RNA editing in plants and its evolution. Annual Review of Genetics 47, 335–352, doi:10.1146/annurev-genet-111212-133519 (2013).

36 Chiu, W.-L. & Sears, B. B. Recombination between chloroplast DNAs does not occur in sexual crosses of *Oenothera*. Mol. Gen. Genet. 198, 525–528 (1985).

37 Greiner, S., Sobanski, J. & Bock, R. Why are most organelle genomes transmitted maternally? Bioessays 37, 80–94, doi:10.1002/bies.201400110 (2015).

38 Simon, M. et al. Genomic conflicts that cause pollen mortality and raise reproductive barriers in *Arabidopsis thaliana*. Genetics 203, 1353–1367, doi:10.1534/genetics.115.183707 (2016).

39 Sobanski, J. et al. Chloroplast competition is controlled by lipid biosynthesis in evening primroses. Proceedings of the National Academy of Sciences 116, 5665–5674, doi:10.1073/pnas.1811661116 (2019).

40 Stubbe, W. Genetische Analyse des Zusammenwirkens von Genom und Plastom bei *Oenothera*. Zeitschrift für Vererbungslehre 90, 288–298 (1959).

41 Stubbe, W. Untersuchungen zur genetischen Analyse des Plastoms von *Oenothera*. Zeitschrift für Botanik 48, 191–218 (1960).

42 Stubbe, W. Die Rolle des Plastoms in der Evolution der Oenotheren. Ber. Dtsch. Bot. Ges. 76, 154–167 (1963).

43 Wasmund, O. Cytogenetische Untersuchung zur Systematik einiger Sippen der Subsektion Euoenothera der Gattung Oenothera (Onagraceae) Erste Staatsprüfung Lehramt für Sekundarstufe II thesis, Heinrich-Heine-Universität, (1980).

44 Burrows, P. A., Sazanov, L. A., Svab, Z., Maliga, P. & Nixon, P. J. Identification of a functional respiratory complex in chloroplasts through analysis of tobacco mutants containing disrupted plastid *ndh* genes. EMBO J. 17, 868–876, doi:10.1093/emboj/17.4.868 (1998).

45 Umate, P. et al. Deletion of *psbM* in tobacco alters the QB site properties and the electron flow within photosystem II. Journal of Biological Chemistry 282, 9758–9767, doi:10.1074/jbc.M608117200 (2007).

46 Schwenkert, S. et al. Role of the low-molecular-weight subunits PetL, PetG, and PetN in assembly, stability, and dimerization of the cytochrome *b*_6_*f* complex in tobacco. Plant Physiology 144, 1924–1935, doi:10.1104/pp.107.100131 (2007).

47 Hager, M., Biehler, K., Illerhaus, J., Ruf, S. & Bock, R. Targeted inactivation of the smallest plastid genome-encoded open reading frame reveals a novel and essential subunit of the cytochrome *bf* complex. EMBO J. 18, 5834–5842, doi:10.1093/emboj/18.21.5834 (1999).

48 Anderson, J. M., Dean Price, G., Soon Chow, W., Hope, A. B. & Badger, M. R. Reduced levels of cytochrome bf complex in transgenic tobacco leads to marked photochemical reduction of the plastoquinone pool, without significant change in acclimation to irradiance. Photosynthesis Research 53, 215–227, doi:10.1023/a:1005856615915 (1997).

49 Schöttler, M. A., Flügel, C., Thiele, W. & Bock, R. Knock-out of the plastid-encoded PetL subunit results in reduced stability and accelerated leaf age-dependent loss of the cytochrome *b_6_f* complex. Journal of Biological Chemistry 282, 976–985, doi:10.1074/jbc.M606436200 (2007).

50 Zupok, A. The psbB operon is a major locus for plastome-genome incompatibility in Oenothera PhD thesis, University of Potsdam, (2015).

51 Zghidi-Abouzid, O., Merendino, L., Buhr, F., Malik Ghulam, M. & Lerbs-Mache, S. Characterization of plastid *psbT* sense and antisense RNAs. Nucleic Acids Research 39, 5379–5387, doi:10.1093/nar/gkr143 (2011).

52 Chevalier, F. et al. Characterization of the *psbH* precursor RNAs reveals a precise endoribonuclease cleavage site in the *psbT/psbH* intergenic region that is dependent on *psbN* gene expression. Plant Molecular Biology 88, 357–367, doi:10.1007/s11103-015-0325-y (2015).

53 Krech, K. et al. Reverse genetics in complex multigene operons by co-transformation of the plastid genome and its application to the open reading frame previously designated *psbN*. The Plant Journal 75, 1062–1074 (2013).

54 Torabi, S. et al. PsbN is required for assembly of the photosystem II reaction center in *Nicotiana tabacum*. Plant Cell 26, 1183–1199 (2014).

55 Bogdanova, V. S. et al. Nuclear-cytoplasmic conflict in pea *(Pisum sativum* L.) is associated with nuclear and plastidic candidate genes encoding acetyl-CoA carboxylase subunits. PLoS ONE 10, e0119835, doi:10.1371/journal.pone.0119835 (2015).

56 Nováková, E. et al. Allelic Diversity of Acetyl Coenzyme A Carboxylase accD/bccp Genes Implicated in Nuclear-Cytoplasmic Conflict in the Wild and Domesticated Pea (Pisum sp.). International Journal of Molecular Sciences 20, 1773 (2019).

57 Sambatti, J. B. M., Ortiz-Barrientos, D., Baack, E. J. & Rieseberg, L. H. Ecological selection maintains cytonuclear incompatibilities in hybridizing sunflowers. Ecology Letters 11, 1082–1091, doi:10.1111/j.1461-0248.2008.01224.x (2008).

58 Chi, W., He, B., Mao, J., Jiang, J. & Zhang, L. Plastid sigma factors: their individual functions and regulation in transcription. Biochimica et Biophysica Acta 1847, 770–778, doi:https://doi.org/10.1016/j.bbabio.2015.01.001 (2015).

59 Noordally, Z. B. et al. Circadian control of chloroplast transcription by a nuclear-encoded timing signal. Science 339, 1316–1319, doi:10.1126/science.1230397 (2013).

60 Shimizu, M. et al. Sigma factor phosphorylation in the photosynthetic control of photosystem stoichiometry. Proceedings of the National Academy of Sciences 107, 10760–10764, doi:10.1073/pnas.0911692107 (2010).

61 Zhang, J., Ruhlman, T. A., Sabir, J., Blazier, J. C. & Jansen, R. K. Coordinated rates of evolution between interacting plastid and nuclear genes in Geraniaceae. Plant Cell 27, 563–573, doi:10.1105/tpc.114.134353 (2015).

62 Kuroda, H. & Sugiura, M. Processing of the 5’-UTR and existence of protein factors that regulate translation of tobacco chloroplast psbN mRNA. Plant Molecular Biology 86, 585–593, doi:10.1007/s11103-014-0248-z (2014).

63 Harte, C. Oenothera – Contributions of a Plant to Biology. 1st edn, (Springer, 1994).

64 Schumacher, E. & Steiner, E. E. Cytological analysis of complex-heterozygotes in populations of *Oenothera grandiflora* (Onagraceae) in Alabama. Plant Systematics and Evolution 184, 77–87 (1993).

65 Schumacher, E., Steiner, E. E. & Stubbe, W. The complex-heterozygotes of *Oenothera grandiflora* L’Her. Bot. Acta 105, 375–381 (1992).

66 Steiner, E. E. & Stubbe, W. A contribution to the population biology of *Oenothera grandiflora* L’Her. American Journal of Botany 71, 1293–1301 (1984).

67 Steiner, E. E. & Stubbe, W. *Oenothera grandiflora* revisited: a new view of its population structure. Bull Torrey Bot Club 113, 406–412 (1986).

68 Stubbe, W. & Raven, P. H. Genetic self-incompatibility in *Oenothera* subsect *Euoenothera*. Science 204, 327, doi:10.1126/science.204.4390.327 (1979).

69 Stubbe, W. & Steiner, E. Inactivation of pollen and other effects of genome-plastome incompatibility in *Oenothera*. Plant Systematics and Evolution 217, 259–277 (1999).

70 Wasmund, O. Cytogenetic investigation on *Oenothera nutans* (Onagraceae). Plant Systematics and Evolution 169, 69–80 (1990).

71 Wasmund, O. & Stubbe, W. Cytogenetic investigations on *Oenothera wolfii* (Onagraceae). Plant Systematics and Evolution 154, 79–88 (1986).

72 Cleland, R. E. Plastid behaviour of the North American Euoenotheras. Planta 57, 699–712 (1962).

73 Stubbe, W. *Oenothera* – An ideal system for studying the interaction of genome and plastome. Plant Mol. Biol. Rep. 7, 245–257 (1989).

74 Schötz, F. Über Plastidenkonkurrenz bei *Oenothera*. Planta 43, 182–240 (1954).

75 Chiu, W.-L., Stubbe, W. & Sears, B. B. Plastid inheritance in *Oenothera:* organelle genome modifies the extent of biparental plastid transmission. Current Genetics 13, 181–189 (1988).

76 Porra, R. J., Thompson, W. A. & Kriedemann, P. E. Determination of accurate extinction coefficients and simultaneous equations for assaying chlorophylls a and b extracted with four different solvents: verification of the concentration of chlorophyll standards by atomic absorption spectroscopy. Biochimica et Biophysica Acta 975, 384–394, doi:http://dx.doi.org/10.1016/S0005-2728(89)80347-0 (1989).

77 Schöttler, M. A., Kirchhoff, H. & Weis, E. The role of plastocyanin in the adjustment of the photosynthetic electron transport to the carbon metabolism in tobacco. Plant Physiology 136, 4265, doi:10.1104/pp.104.052324 (2004).

78 Schöttler, M. A., Flügel, C., Thiele, W. & Bock, R. The plastome-encoded PsaJ subunit is required for efficient photosystem I excitation, but not for plastocyanin oxidation in tobacco. Biochem J. 403, 251–260 (2007).

79 Kirchhoff, H., Mukherjee, U. & Galla, H. J. Molecular architecture of the thylakoid membrane: lipid diffusion space for plastoquinone. Biochemistry 41, 4872–4882, doi:10.1021/bi011650y (2002).

80 Rott, M. et al. ATP synthase repression in tobacco restricts photosynthetic electron transport, CO_2_ assimilation, and plant growth by overacidification of the thylakoid lumen. Plant Cell 23, 304–321, doi:10.1105/tpc.110.079111 (2011).

81 Schwenkert, S. et al. PsbI affects the stability, function, and phosphorylation patterns of photosystem II assemblies in tobacco. J Biol Chem 281, 34227–34238, doi:10.1074/jbc.M604888200 (2006).

82 Ossenbühl, F. et al. Efficient assembly of photosystem II in *Chlamydomonas reinhardtii* requires Alb3.1p, a homolog of *Arabidopsis* ALBINO3. Plant Cell 16, 1790–1800, doi:10.1105/tpc.023226 (2004).

83 Massouh, A. et al. Spontaneous chloroplast mutants mostly occur by replication slippage and show a biased pattern in the plastome of *Oenothera*. Plant Cell 28, 911–929, doi:10.1105/tpc.15.00879 (2016).

84 Lehwark, P. & Greiner, S. GB2sequin – a file converter preparing custom GenBank files for database submission. Genomics 111, 759–761, doi:https://doi.org/10.1016/j.ygeno.2018.05.003 (2019).

85 Mesquite: a modular system for evolutionary analysis v. 3.40 (2018).

86 Leebens-Mack, J. H. et al. One thousand plant transcriptomes and the phylogenomics of green plants. Nature 574, 679–685, doi:10.1038/s41586-019-1693-2 (2019).

87 Kühn, K., Weihe, A. & Börner, T. Multiple promoters are a common feature of mitochondrial genes in Arabidopsis. Nucleic acids research 33, 337–346, doi:10.1093/nar/gki179 (2005).

## Supplementary References

1 Schöttler, M. A. & Tóth, S. Z. Photosynthetic complex stoichiometry dynamics in higher plants: environmental acclimation and photosynthetic flux control. Front. Plant Sci. 5, 188–188, doi:10.3389/fpls.2014.00188 (2014).

2 Krause, G. H. & Weis, E. Chlorophyll fluorescence and photosynthesis: the Basics. Annu. Rev. Plant Physiol. Plant Mol. Biol. 42, 313–349, doi:10.1146/annurev.pp.42.060191.001525 (1991).

3 Albus, C. A. et al. Y3IP1, a nucleus-encoded thylakoid protein, cooperates with the plastid-encoded Ycf3 protein in photosystem I assembly of tobacco and *Arabidopsis*. Plant Cell 22, 2838–2855, doi:10.1105/tpc.110.073908 (2010).

4 Krech, K. et al. The plastid genome-encoded Ycf4 protein functions as a nonessential assembly factor for photosystem I in higher plants. Plant Physiology 159, 579–591, doi:10.1104/pp.112.196642 (2012).

5 Rott, M. et al. ATP synthase repression in tobacco restricts photosynthetic electron transport, CO2 assimilation, and plant growth by overacidification of the thylakoid lumen. Plant Cell 23, 304–321, doi:10.1105/tpc.110.079111 (2011).

6 Cruz, J. A., Sacksteder, C. A., Kanazawa, A. & Kramer, D. M. Contribution of electric field *(Δψ)* to steady-state transthylakoid proton motive force *(pmf) in vitro* and *in vivo.* Control of *pmf* parsing into Δψ and ΔpH by Ionic strength. Biochemistry 40, 1226–1237, doi:10.1021/bi0018741 (2001).

7 Westhoff, P. & Herrmann, R. G. Complex RNA maturation in chloroplasts. The *psbB* operon from spinach. European Journal of Biochemistry 171, 551–564 (1988).

8 Stubbe, W. Die Rolle des Plastoms in der Evolution der Oenotheren. Ber. Dtsch. Bot. Ges. 76, 154–167 (1963).

9 Drillisch, M. Vergleichende Untersuchungen an “A-Genotypen” von Oenothera PhD thesis, Heinrich-Heine-Universität, (1975).

10 Dietrich, W., Wagner, W. L. & Raven, P. H. Systematics of Oenothera section Oenothera subsection Oenothera (Onagraceae). 1st edn, (The American Society of Plant Taxonomists, 1997).

11 Massouh, A. et al. Spontaneous chloroplast mutants mostly occur by replication slippage and show a biased pattern in the plastome of *Oenothera*. Plant Cell 28, 911–929, doi:10.1105/tpc.15.00879 (2016).

12 Stubbe, W. Untersuchungen zur genetischen Analyse des Plastoms von *Oenothera*. Zeitschrift für Botanik 48, 191–218 (1960).

13 Stubbe, W. Genetische Analyse des Zusammenwirkens von Genom und Plastom bei *Oenothera*. Zeitschrift für Vererbungslehre 90, 288–298 (1959).

14 Cleland, R. E. Plastid behaviour of the North American Euoenotheras. Planta 57, 699–712 (1962).

15 Sobanski, J. et al. Chloroplast competition is controlled by lipid biosynthesis in evening primroses. Proceedings of the National Academy of Sciences 116, 5665–5674, doi:10.1073/pnas.1811661116 (2019).

16 Kozul, D. Systematic identification of loci determining chloroplast and nuclear genome incompatibilities in the evening primrose (Oenothera) PhD thesis, University of Potsdam, (2019).

17 Hupfer, H. et al. Complete nucleotide sequence of the *Oenothera elata* plastid chromosome, representing plastome I of the five distinguishable *Euoenothera* plastomes. Mol. Gen. Genet. 263, 581–585 (2000).

18 Greiner, S. et al. The complete nucleotide sequences of the five genetically distinct plastid genomes of *Oenothera*, subsection *Oenothera:* I. Sequence evaluation and plastome evolution. Nucleic Acids Research 36, 2366–2378, doi:10.1093/nar/gkn081 (2008).

19 Greiner, S. et al. The complete nucleotide sequences of the 5 genetically distinct plastid genomes of *Oenothera*, subsection *Oenothera:* II. A microevolutionary view using bioinformatics and formal genetic data. Molecular Biology and Evolution 25, 2019–2030, doi:10.1093/molbev/msn149 (2008).

20 Wasmund, O. Cytogenetische Untersuchung zur Systematik einiger Sippen der Subsektion Euoenothera der Gattung Oenothera (Onagraceae) Erste Staatsprüfung Lehramt für Sekundarstufe II thesis, Heinrich-Heine-Universität, (1980).

21 Wasmund, O. & Stubbe, W. Cytogenetic investigations on *Oenothera wolfii* (Onagraceae). Plant Systematics and Evolution 154, 79–88 (1986).

22 Schumacher, E., Steiner, E. E. & Stubbe, W. The complex-heterozygotes of *Oenothera grandiflora* L’Her. Bot. Acta 105, 375–381 (1992).

23 Hupfer, H. Vergleichende Sequenzanalyse der fünf Grundplastome der Sektion Oenothera (Gattung Oenothera) – Analyse des Cytochrom-Komplexes PhD thesis, Ludwig-Maximilians-University, (2002).

24 Keuthe, M. Identifizierung und Charakterisierung von Suppressormutanten einer Plastom-Genom-Inkompatibilität in Oenothera PhD thesis, University of Potsdam, (2013).

25 Munz, P. A. The *Oenothera hookeri* group. El Aliso 2, 1–47 (1949).

26 Steiner, E. E. A cytogenetic study of certain races of *Oenothera elata*. Bull Torrey Bot Club 82, 292–297 (1955).

27 Steiner, E. E. Phylogenetic relationships of certain races of *Euoenothera* from Mexico and Guatemala. Evolution 5, 265–272 (1951).

28 Cleland, R. E. The reduction divisions in the pollen mother cells of *Oenothera franciscana*. American Journal of Botany 9, 391–413, doi:10.2307/2435274 (1922).

29 Bartlett, H. H. Systematic studies on *Oenothera:* IV. *Oe. franciscana* and *Oe. venusta*, spp. novv. Rhodora 16, 33–37 (1914).

30 de Vries, H. Gruppenweise Artbildung – Unter spezieller Berücksichtigung der Gattung Oenothera. (Gebrüder Borntraeger, 1913).

31 Cleland, R. E. Cyto-taxonomic studies on certain Oenotheras from California. Proc. Am. Philos. Soc. 75, 339–429 (1935).

32 Baerecke, M. Zur Genetik und Cytologie von *Oenothera ammophila* Focke, *Bauri* Boedijn, *Beckeri* Renner, *parviflora* L., *rubricaulis* Klebahn, *silesiaca* Renner. Flora 138, 57–92 (1944).

33 Renner, O. Wilde Oenotheren in Norddeutschland. Flora 31, 182–226 (1937).

34 Cleland, R. E. & Hammond, B. L. in Studies in Oenothera cytogenetics and phylogeny Vol. 16 (ed Ralph E. Cleland) 10–72 (Indiana University Publications. Science Series, 1950).

35 Cleland, R. E. Species relationships in *Onagra*. Proc. Am. Philos. Soc. 77, 477–542 (1937).

36 Renner, O. & Hirmer, U. Zur Kenntnis von *Oenothera:* I. Über *Oe. conferta* n. sp. II. Über künstliche Polyploidie. Biologisches Zentralblatt 75, 513–531 (1956).

37 Rossmann, G. Analyse der *Oenothera coronifera* Renner. Flora 153, 451–468 (1963).

38 Renner, O. Europäische Wildarten von *Oenothera:* III. Planta 47, 219–254 (1956).

39 Renner, O. Versuche über die gametische Konstitution der Önotheren. Z. indukt. Abstamm. Vererbungsl. 18, 121–294 (1917).

40 Klebahn, H. Formen, Mutationen, und Kreuzungen bei einigen Oenotheren aus der Lüneburger Heide. Jahrbuch der Hamburger Wissenschaftlichen Anstalten 31, 1–64 (1914).

41 Klebahn, H. Weitere Beobachtungen über Oenotheren aus Nordwestdeutschland. Z. indukt. Abstamm. Vererbungsl. 39, 8–30, doi:10.1007/BF01961518 (1925).

42 Renner, O. Kurze Mitteilung über *Oenothera:* V. Zur Kenntnis von *O. silesiaca* n. sp., *parviflora* L., *ammophila* Focke, *rubricaulis* Kleb. Flora 36, 325–335 (1943).

43 de Vries, H. Mutations of *Oenothera suaveolens* Desf. Genetics 3, 1–26 (1918).

44 Stubbe, W. Genetische und zytologische Untersuchungen an verschiedenen Sippen von *Oenothera suaveolens*. Z. indukt. Abstamm. Vererbungsl. 85, 180–209 (1953).

45 Renner, O. Europäische Wildarten von *Oenothera:* II. Ber. Dtsch. Bot. Ges. 63, 129–138 (1950).

46 Steiner, E. Phylogenetic studies in *Euoenothera*. Evolution 6, 69–80 (1952).

47 Cleland, R. E. The evolution of the North American Oenotheras of the *“biennis”* group. Planta 51, 378–398 (1958).

48 Heribert-Nilsson, N. Die Variabilität der *Oenothera Lamarckiana* und das Problem der Mutation. Zeitschrift für induktive Abstammungs-und Vererbungslehre 8, 89–231 (1912).

49 Steiner, E. E. & Stubbe, W. A contribution to the population biology of *Oenothera grandiflora* L’Her. American Journal of Botany 71, 1293–1301 (1984).

50 Hoeppener, E. & Renner, O. Genetische und zytologische Oenotherenstudien: I. Zur Kenntnis der *Oenothera ammophila* Focke. Z. indukt. Abstamm. Vererbungsl. 49, 1–25 (1929).

51 Boedijn, K. Die systematische Gruppierung der Arten von Oenothera. Z. indukt. Abstamm. Vererbungsl. 32, 354–362, doi:10.1007/BF01816761 (1924).

52 Geckler, H. in Studies in Oenothera cytogenetics and phylogeny Vol. 16 (ed Ralph E. Cleland) 160–217 (Indiana University Publications. Science Series, 1950).

53 Renner, O. Über *Oenothera atrovirens* Sh. et Bartl. und über somatische Konversion im Erbgang des cruciata-Merkmals der Oenotheren. Z. indukt. Abstamm. Vererbungsl. 74, 91124 (1938).

54 Davis, B. M. Hybrids of *Oenothera biennis* and *Oenothera franciscana* in the first and second generations. Genetics 1, 197–251 (1916).

55 Renner, O. Über die Entstehung homozygotischer Formen aus kompelx-heterozygotischen Oenotheren. Flora 135, 201–238 (1941).

56 Cleland, R. E. Oenothera – Cytogenetics and Evolution. 1st edn, (Academic Press Inc., 1972).

57 Bartlett, H. H. An account of the cruciate-flowered Oenotheras of the subgenus *Onagra*. American Journal of Botany 1, 226–243, doi:10.2307/2435255 (1914).

58 Stubbe, W. *Oenothera* – An ideal system for studying the interaction of genome and plastome. Plant Mol. Biol. Rep. 7, 245–257 (1989).

